# The cytoplasm of living cells can sustain transient and steady intracellular pressure gradients

**DOI:** 10.1101/2024.04.22.590593

**Authors:** Majid Malboubi, Mohammad Hadi Esteki, Malti B. Vaghela, Lulu I T. Korsak, Ryan J. Petrie, Emad Moeendarbary, Guillaume Charras

## Abstract

Understanding the physical basis of cellular shape change in response to both internal and external mechanical stresses requires characterisation of cytoplasmic rheology. At subsecond time-scales and micron length-scales, cells behave as fluid-filled sponges in which shape changes necessitate intracellular fluid redistribution. However, whether these cytoplasmic poroelastic properties play an important role in cellular mechanical response over length-scales and time-scales relevant to cell physiology remains unclear. Here, we investigated whether and how a localised deformation of the cell surface gives rise to transient intracellular flows spanning several microns and lasting seconds. Next, we showed that pressure gradients induced in the cytoplasm can be sustained over several minutes. We found that stable pressure gradients can arise from the combination of cortical tension, cytoplasmic poroelasticity and water flows across the membrane. Overall our data indicate that intracellular cytosolic flows and pressure gradients may play a much greater role than currently appreciated, acting over time- and length-scales relevant to mechanotransduction and cell migration, signifying that poroelastic properties need to be accounted for in models of the cell.

**Significance statement:** Understanding how cells change shape dynamically under the influence of external and internal forces requires characterisation of the mechanical response of the cytoplasm, the viscous material that fills their interior. The cytoplasm consists of a porous solid phase bathed in a fluid, the cytosol. As the cytoplasm is incompressible, any cellular shape change necessitates redistribution of the cytosol within the cell and its flow rate sets the time-scale for deformation. How the cytoplasmic mechanical response affects cell physiology remains poorly understood. We show that the unique physical properties of the cytoplasm allow cells to sustain cellular-scale pressure gradients over minute time-scales. As a consequence, pressure-driven mechanisms may play a much greater role in cell physiology than currently appreciated.

## Introduction

The rheology of cells determines their response to internal and external mechanical stimuli encountered during normal physiological function. Since the cytoplasm forms the largest part of the cell by volume, its rheological properties are key to understanding cellular shape change in response to mechanical stress. The cytoplasm can be described as a biphasic material that consists of a porous solid phase bathed in an interstitial fluid^1^. In such a material, stress relaxation arises from fluid flow through the pores of the solid phase as a result of spatial gradients in the applied stress field^2^. By studying stress relaxation in micro-indentation experiments combined with perturbations including volumetric deformations together with chemical and genetic treatments, previous work has shown that, at sub-second time-scales and micron length-scales, the cytoplasm behaves as a poroelastic material^1,3^. However, the contribution of cytoplasmic poroelastic properties to the mechanical response of cells has not been characterized on the tens of micron length-scale and second-to-minute time-scale relevant to cell physiology. Thus, it is unclear if poroelastic properties must be accounted for in models of cell mechanics.

During mechanotransduction, cells sense external mechanical stresses and translate them into biochemical signals^4^. The intracellular fluid flows and pressure gradients elicited in poroelastic materials in response to local application of stress may represent a stimulus for detection. While deformations are applied at the cellular scale, most mechanosensory processes act at the molecular scale through opening of ion channels and unfolding of proteins. Thus, the spatial extent of the stress and strain fields will determine what proportion of the cell is likely to respond to a stimulus and the temporal evolution of these stress and strain fields will determine the duration over which mechanosensory processes will be stimulated. Understanding how stresses equilibrate in the cytoplasm in response to local application of force is necessary to better understand the physical parameters detected by intracellular mechanotransductory pathways. However, direct characterisation of the spatiotemporal deformation induced by sudden local application of force is currently lacking.

Migrating cells often adopt a polarised morphology with a gradient of non-muscle myosin II (NMII) protein increasing towards the rear^5–7^ or the front^8,9^, depending on cell type. Given the poroelastic nature of cytoplasm, theoretical considerations predict that such cortical myosin gradients should result in intracellular pressure gradients driving intracellular fluid flows in the direction opposite to the gradient of NMII^10,11^. Intracellular flows have been proposed to participate in protrusion formation and migration but have only been indirectly observed in cells^12–17^. These flows may play a particularly important role during migration in confined environments with low adhesion, where most cell types adopt an amoeboid morphology and extend pressure-driven protrusions at their front^5–7,18–20^. Although significant evidence points towards a role for myosin-generated pressure gradients in migration^9^, it is unclear if intracellular pressure gradients can be sustained over the minute-long time-scales involved in migration.

Here, we used a combination of cell physiology experiments, high-resolution nanoparticle tracking, and finite element (FE) simulations to study how the combination of membrane permeability and cytoplasmic poroelastic properties can lead to steady-state gradients in intracellular pressure. We investigate the dynamics of cellular stress relaxation in response to external and internal mechanical stresses to determine if the poroelastic nature of cytoplasm is relevant to cell physiology on tens of microns length-scales and minute time- scales. We observe a cell-scale mechanical response following application of a local deformation of the cell surface and show that it is due to transient intracellular fluid flows elicited in poroelastic materials. We then reveal experimentally that intracellular pressure gradients lasting several minutes can be sustained in the cytoplasm, signifying that pressure gradients may play a role in migration.

## Results and discussion

### Whole-cell mechanical equilibration in response to localised deformation necessitates several seconds

Cells are often subjected to localised mechanical forces applied at high strain rates^21–23^. In poroelastic materials, rapid application of localised stress pressurises the interstitial fluid. Stress relaxes by flow of water out of the deformed region through the pores of the solid phase. As a consequence, the rate of mechanical equilibration is set by the poroelastic diffusion constant *D_p_* that depends on the elasticity 𝐸 of the solid phase, the viscosity *η* of the fluid phase, and the size 𝜉 of the pores through which the interstitial fluid can permeate: *D_p_*∼𝐸𝜉^2^/*η* ^24^. The time-scale of water flow out of the deformed region is 𝑡_𝑝_∼𝐿^2^/*D_p_*, where 𝐿 is a length-scale associated with deformation. Previous work has shown that local force application to a cell can result in global changes in the height of the cell surface and revealed the presence of two regimes: one fast due to stress propagation in the cytoskeleton and the other slow, whose origin was unclear^25^. As previous work has shown that the cytoplasm behaves as a poroelastic material^1^, we hypothesized that the slow response taking place of over tens of microns length-scale and seconds time-scale reflects the poroelastic nature of the cytoplasm.

To examine the role of poroelasticity in the mechanical response of cells, we locally deformed the cell surface with the tip of an AFM cantilever. In our experiments, the deformation had an estimated characteristic length 𝐿∼3 𝜇𝑚 (𝐿∼√𝑑𝛿 with d*∼4μm* the diameter of the indentation and δ∼2 *μm* the indentation depth) applied with a rise time 𝑡_𝑟_∼100𝑚𝑠 (**Fig 1A**, **B**). In Hela cells, previous work has reported *D_p_*∼40𝜇𝑚^2^/𝑠 ^26^, leading to an estimate of the characteristic time of fluid efflux 𝑡_𝑝_∼200 𝑚𝑠 . As 𝑡_𝑝_ is larger than the rise time 𝑡_𝑟_ , poroelasticity may contribute to mechanical relaxation.

**Figure 1:**
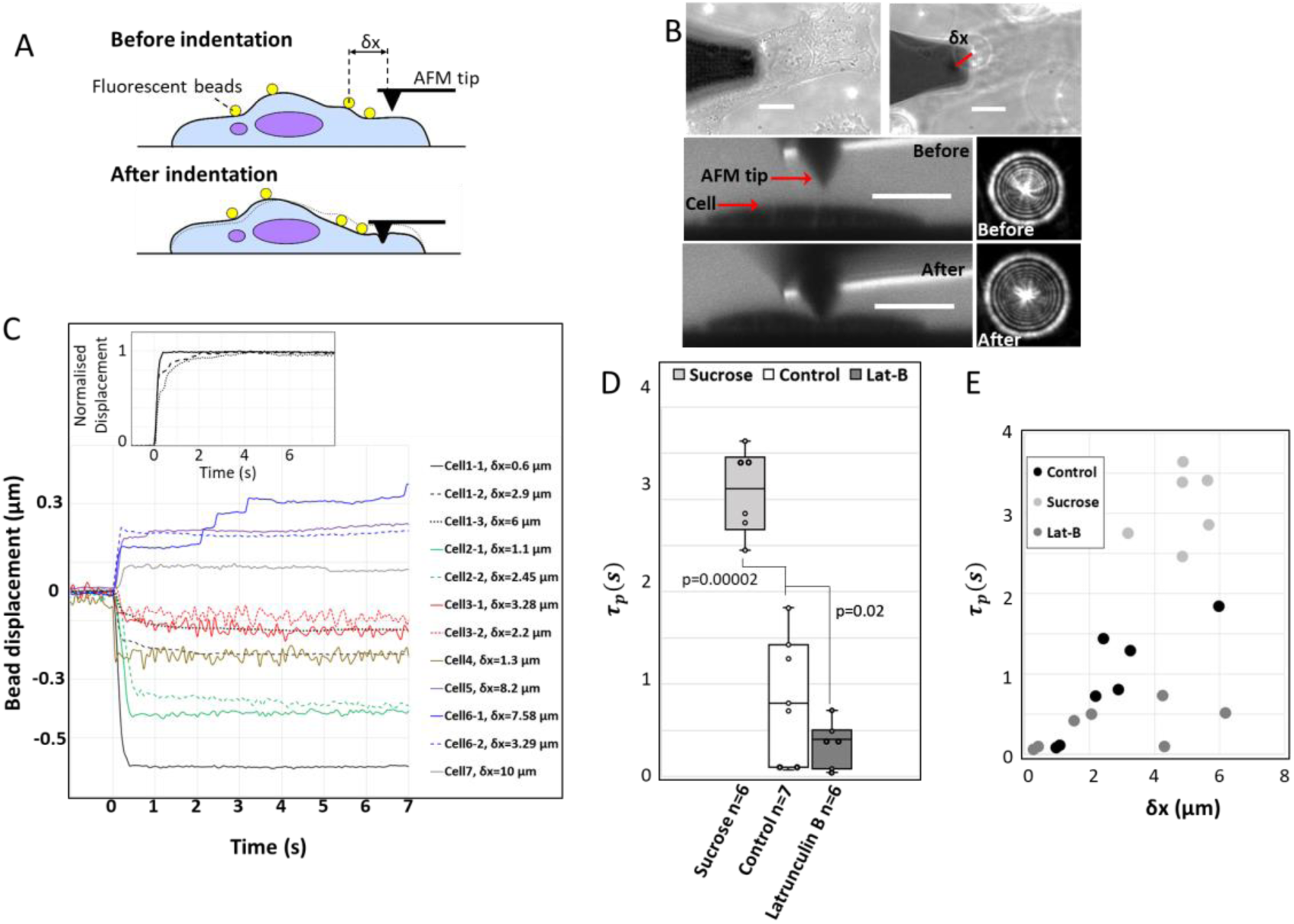
Local application of stress gives rise to cell-scale intracellular flow. **A.** Schematic diagram of the experiment. Collagen-coated fluorescent beads are bound to the cell surface. An AFM cantilever is brought into contact with one side of the cell and bead movement in the z-direction is imaged over time. **B.** Representative image showing a combined phase contrast and fluorescence image of the beads on the cell. Top panels: left: The AFM cantilever appears as a dark shadow on the left of the image. The bead is visualized by fluorescence. When the plane of focus is moved higher than the cell, a halo of fluorescence centered on the bead appears (middle panel, right). The diameter of the halo of the bead reports on the distance between the bead and the plane of focus (see SI, **Fig S1A**). Variations in halo radius indicate changes in height caused by indentation. The distance between the bead and the AFM tip is indicated by a red line. Middle panels: left: Profile of a cell before indentation. A cell- impermeable fluorescent indicator has been added to the medium and the cell appears dark. The AFM cantilever was imaged by reflectance and appears bright. Right: Representative image of the halo before indentation. Bottom panels: left: Profile of the same cell as in the middle panel during indentation by an AFM cantilever. Right: Halo of the same bead as in the middle panel during indentation. Scale bar= 10μm. **C.** Change in bead height as a function of time for a total of 12 beads on 7 cells. Beads from the same cell appear in the same colour. Inset shows normalized displacement of three beads on the same cell located at different distances from the AFM tip. This highlights the slower response in the second phase for more distant beads. The color code in the inset is the same as the main figure. **D.** Characteristic relaxation time *τ_p_* of the second phase for control cells (n=7 cells) and cells treated with sucrose (n=6 cells) and latrunculin (n=7 cells). In the box plot, the black line is the median, the bottom and top edges of the box indicate the 25th and 75^th^ percentiles, respectively. The whiskers extend to the most extreme data points that are not outliers. Data points appear as black dots. **E**. Characteristic relaxation time *τ_p_* as a function of distance from the AFM tip for control cells (black), sucrose (light grey), and latrunculin (dark grey).

To detect the global response of the cell surface to local force application with high spatial accuracy, we employed defocusing microscopy^3,25^. In this technique, collagen-coated fluorescent beads are tethered to the cell surface. The plane of focus is chosen such that the beads are deliberately out of focus and display multiple diffraction rings about their centre (**Fig S1A**). Bead vertical displacement is monitored with nanometre precision by measuring the temporal evolution of the diameter of the outer diffraction ring in response to the deformation of the cell surface induced by AFM (**Fig S1A, B**). At steady-state, beads closer than ∼6𝜇𝑚 to the AFM tip moved downwards while beads further away moved upwards, as expected for an elastic material subjected to indentation and as previously observed^25^ (**Fig 1C**). The amplitude of the steady-state displacement decreased with distance between the bead and the AFM tip (**Fig 1C**, **Fig S3A**). The temporal evolution of the bead movement displayed a biphasic response at all positions throughout the cell (**Fig 1C**). The first phase consisted in a fast movement of all the beads independent of their distance from the AFM tip over a time-scale shorter than ∼0.3 s, possibly due to cell surface tension (**Fig 1C inset, Fig S2**). In the second phase, the beads relaxed to their final displacement over a characteristic time scale 𝜏_𝑝_ that increased with increasing distance between the AFM tip and the bead (**Fig S2, Fig 1C**, **Fig 1D**, **Fig 1E**, **Methods**). The overall relaxation of the second phase lasted up to ∼5s. Thus, local application of external force leads to a whole-cell mechanical equilibration lasting several seconds, consistent with previous work^25^.

When stress is applied to a poroelastic material, relaxation takes place through flow of fluid out of the pressurised region at a rate that depends on the hydraulic permeability of the cytoplasm and hence its pore size. If the slow relaxation observed in cells is indeed due to poroelastic effects driving cellular-scale intracellular fluid flows, changes in the hydraulic pore size ξ should affect the duration of the second phase of bead movement but not the first phase. Pore size can be decreased by increasing the osmolarity of the medium^1,3^. Increasing osmolarity by 300mOsm led to a ∼3.5-fold increase in the median characteristic equilibration time 𝜏_𝑝_ of the slow phase (**Fig 1D**). The amplitude of the first phase of bead movement was unaffected (**Fig S3B**) and we were unable to determine changes in the duration of the first phase because of its short duration. Conversely, when we treated cells with latrunculin, 𝜏_𝑝_ decreased (**Fig 1D**), consistent with previous reports showing that *D_p_* increases with latrunculin treatment ^1^. Surprisingly, the amplitude of bead movement was not affected (**Fig S3**). Thus, the second-long global changes in cell surface height that occur in response to local application of stress are qualitatively consistent with a poroelastic cytoplasm.

### Cytoplasmic flows induced by microinjection follow Darcy’s law

In porous media, pressure imbalances are dissipated by interstitial fluid flows. Though intracellular fluid flows have been inferred to be driven by internal stresses during cell motility^10–12,14,27^ or external stresses during stress relaxation^25,26^, a direct evaluation of the relationship between the pressure gradient 𝛁𝑃_𝑓_ and the velocity of intracellular fluid flow 𝒗 is lacking in cells. In classic porous media, the two are linked by Darcy’s law:

(1) 𝒗 = −𝑘 𝛁𝑃_𝑓_ with *k* the hydraulic permeability of the medium.

To characterise intracellular flows in response to pressure gradients, we created a fluidic link between a micropipette and an interphase cell using techniques developed for electrophysiology (whole-cell recording) and applied a step pressure increase to the micropipette (**Fig 2A**, **Fig S4A, SI methods**). In this configuration, the applied pressure results in fluid injection into the cell, causing the solid phase of the cytoplasm to expand and the height of the cell surface to increase as the fluid progressively permeates through the solid phase of the cytoplasm. In our experiments, we recorded bead displacement using defocusing microscopy for beads at different distances away from the micropipette (**Fig 2B-2C**). After application of a pressure step, beads moved upwards with an amplitude that decreased with increasing distance from the micropipette for a given time (**Fig 2D**) and with a time lag that increased linearly with the distance between the bead and the micropipette (see Methods and **Fig 2E**), indicating an approximately constant velocity of the propagation front. When fluid injection was continued over longer time-scales (>5s), cellular volume regulation mechanisms^28^ could not compensate for the volume increase and the membrane delaminated from the cortex forming large blebs before the cell eventually lysed.

**Figure 2:**
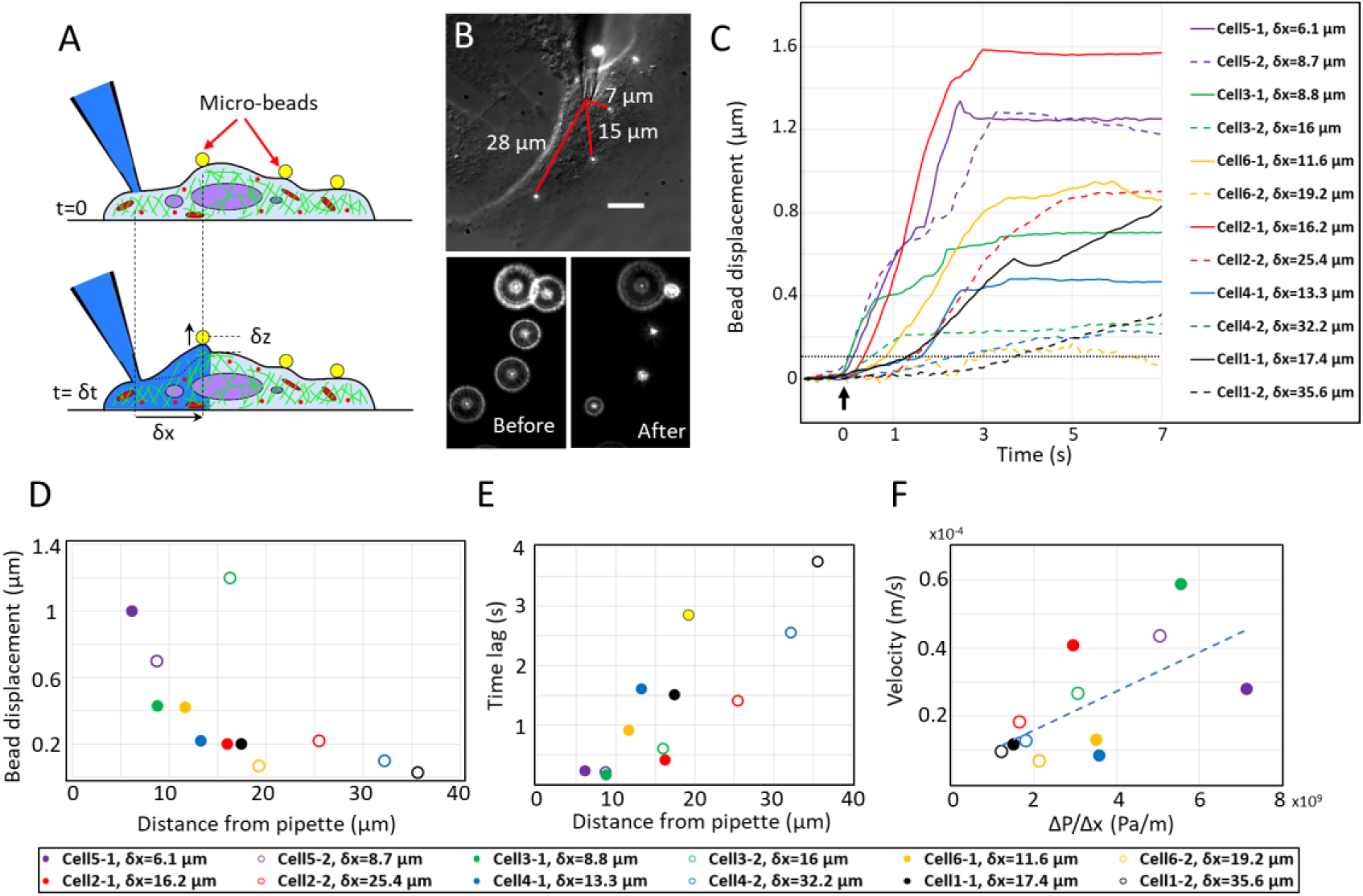
Intracellular fluid propagation in response to pressure gradients. **A.** Schematic diagram of the experiment. Collagen-coated fluorescent beads (yellow) are bound to the surface of a cell in fluidic communication with a micropipette. At time t=0s, pressure is increased within the pipette leading to injection of fluid into the cell (dark blue). After a time lag δt, the fluid propagation front reaches a bead in δx resulting in an increase of its height by δz. **B.** Top: Representative image showing a combined DIC and fluorescence image of a cell. The micropipette appears at the top of the image and a fluidic connection is established. The distance from the pipette tip to each bead is indicated by red lines. Scale bar=10μm. Bottom: Defocused image of the fluorescent beads tethered to the cell surface before (left) and after (right) propagation of the fluid flow through the cell. **C.** Temporal evolution of height for beads situated at different distances from the pipette tip. Data from N=12 beads from n=6 cells. The distance of each bead is listed in the inset. Beads from the same cell are in the same colour. The time at which microinjection starts is t=0 s (indicated by the arrow). The beads respond to injection with a time lag that increases with increasing bead-pipette distance. The dashed black line corresponds to a 0.1 𝜇𝑚 displacement of the beads from their initial position. This threshold is used to calculate the time lag δt between injection and bead response (see SI methods). **D.** Bead displacement as a function of distance from the pipette after t=2s. **E.** Time lag δt as a function of distance from the pipette. **F.** Velocity as a function of estimated pressure gradient. **(D-F)** Beads from the same cell appear as solid or open markers of the same colour. Colour code is indicated on the right of panel F.

In our experiments, the duration of microinjection (∼2s) is short compared to the characteristic time-scale of water flows across the cell membrane (∼20-100s)^29^. Therefore, we can neglect fluid losses through the membrane and we expect flows to follow Darcy’s law in response to microinjection. As the velocity of Darcy flows depends on the hydraulic permeability 𝑘 and the pressure gradient 𝛁𝑃_𝑓_, we plotted the estimated velocity 𝑣∼ ^Δ𝑥^⁄_𝛿𝑡_ as a function of an estimate of the pressure gradient Δ𝑃/Δ𝑥, with Δ𝑥 the position of the bead relative to the micropipette and 𝛿𝑡 the time lag of its movement relative to the onset of injection (see SI for estimation of injection pressure). This revealed a linear dependency with a slope 𝑘∼1.25 10^-13^ m^2^/(Pa.s) (**Fig 2F**, r^2^=0.70), close to previous estimates^1,30^. We then repeated these experiments in metaphase cells in which we could precisely monitor diameter evolution after pressure application (**Fig 3A**). This revealed a strong dependency of the rate of change in diameter *d* and the time-lag 𝛿𝑡 (the onset of diameter increase) with applied pressure (**Fig. 3B-D**). When we plotted an estimate of the velocity 𝑣∼ ^𝑑^⁄_𝛿𝑡_ as a function of an estimate of the pressure gradient 𝛁𝑃_𝑓_∼Δ𝑃/𝑑, this graph again revealed a linear relationship, as expected from Darcy’s law, and yielded an estimate of 𝑘∼5 10^-14^ m^2^/(Pa.s) (**Fig 3E**).

**Figure 3:**
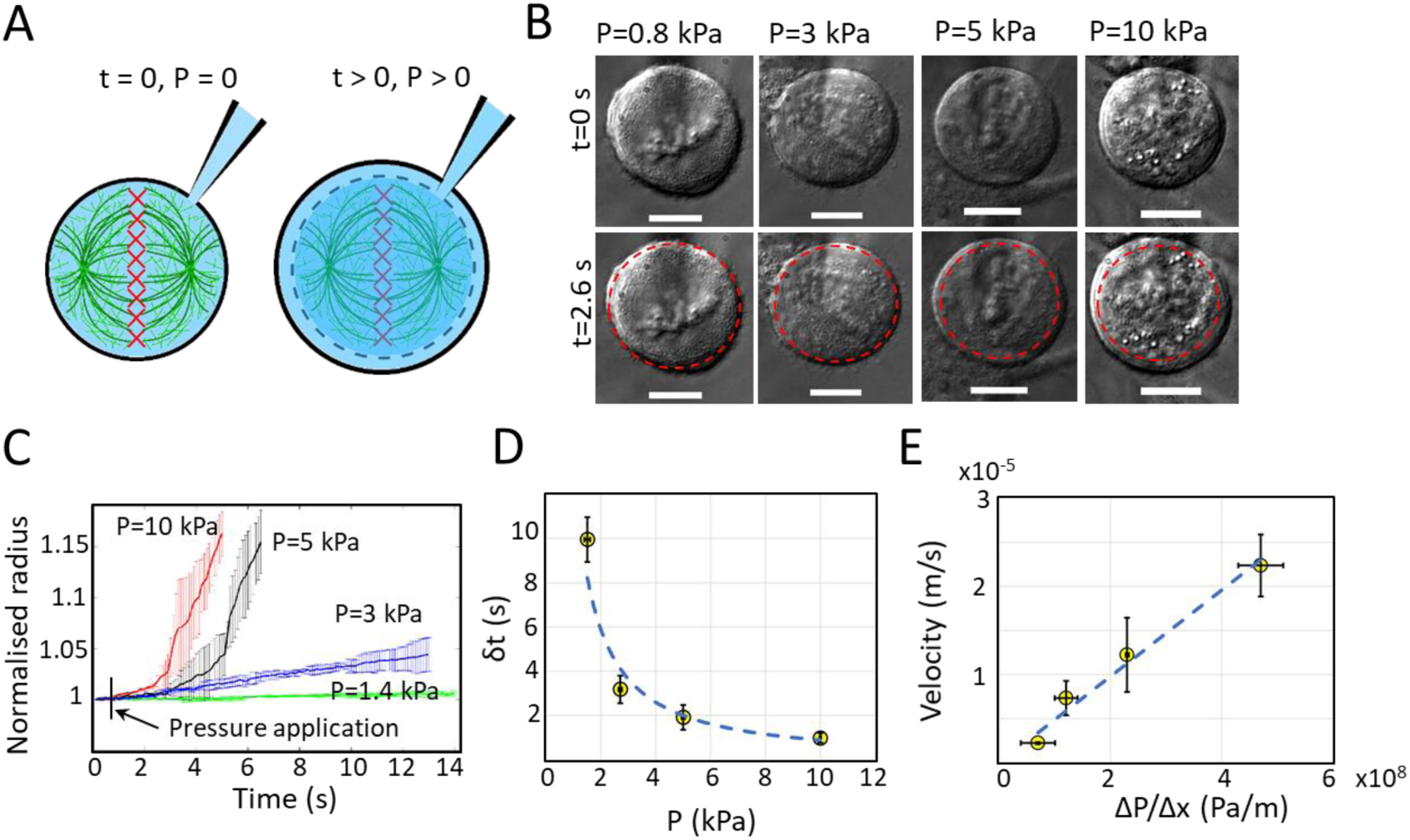
The cytoplasm of metaphase cells displays a porous behavior. **A.** Schematic diagram of the experiment examining the change in diameter of metaphase HeLa cells due to application of a step increase in pressure through the micropipette. To detect the arrival of fluid flow at the cell periphery following application of a step pressure in the micropipette, we monitored changes in cell diameter. An increase in cell diameter was detectable after a time delay *δt* that depended on the amplitude of the pressure step. **B.** DIC images of representative experiments. The step change in pressure was applied at t=0s. Top: equatorial plane of cells blocked in metaphase at t=0s. The micropipette tip is out of the plane of focus. Bottom: cells at t=2.6s after pressure application. The initial diameter is indicated by the red dashed circle. Scale bar=10µm. **C.** Temporal evolution of the relative diameter of metaphase HeLa cells subjected to different pressure steps. Error bars represent standard deviations. N=3 cells per pressure. The timing of pressure application is indicated by a vertical black line. The pressure corresponding to each curve is indicated on the graph. **D.** Time lag δt between pressure application and the onset of diameter increase as a function of the amplitude of the pressure step. The dashed line indicates a hyperbolic fit. Whiskers indicate the standard deviation. n=3 cells per pressure. **E.** Velocity as a function of estimated pressure gradient. The dashed line indicates a linear fit. Whiskers indicate the standard deviation. N=3 experiments per data point.

Together, these experiments show that pressure gradients lead to intracellular flows whose propagation follows Darcy’s law with a hydraulic permeability consistent with those estimated by other experimental approaches ^1,3,30^. Counterintuitively, the hydraulic permeability in interphase cells was higher than in metaphase cells despite cell volume increasing in mitosis. This may be due to the profound remodelling of cytoplasmic organisation accompanying this stage of the cell cycle.

### Global cellular response to microindendation is compatible with a poroelastic cytoplasm

Next, we sought to gain insight into how the global cellular response to local indentation might arise from interplay of cellular poroelastic properties with other cellular mechanical properties. Previous work has hypothesised that local application of mechanical stress leads to a localised outflow of interstitial fluid from the deformed region that can then propagate through the cell^1,25^. If vertical displacements in indentation experiments were just due to fluid propagation, we would expect that the onset of displacement would occur with a time lag that depends on distance as in the microinjection experiments (**Fig 2C-D**). Yet, the first phase of displacement shows no time lag and vertical displacements of the cell surface are observed far from the point of indentation (**Fig 1C-inset**). This indicates that poroelastic properties alone cannot explain the first phase of displacement.

One potential mechanical origin for the first phase may be the submembranous cortex whose mechanics differ markedly from the cytoplasm^31^ and which generates a surface tension 𝛾 through myosin contractility^32^. The relative importance of surface tension in the cortex compared to elastic restoring forces in the cytoplasm can be grasped from the length scale 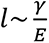 with E the elasticity of the cytoplasm. For length-scales larger than 𝑙, the effect of surface tension is negligible and elastic restoring forces are dominant. Using characteristic values of 𝛾∼1 𝑚𝑁/𝑚 ^33^ and 𝐸_cytoplasm_∼100 Pa ^26^, we obtain a length scale of ∼10 μm, consistent with our experiments (**Fig 1D**). Furthermore, this scaling may explain why latrunculin treatment does not perturb the amplitude of bead movement in the first phase (**Fig S3B**). Indeed, latrunculin affects both the surface tension and the elastic modulus, potentially leading to compensation.

To determine if surface tension is important in setting the scale of the movement of the cell surface, we examined deformation of the membrane induced by indentation of the cell by a sharp AFM tip using mitotic cells to isolate the role of the cortex from cell shape and adhesion (**Fig S5**). The surface profile was imaged before and during a 2 µm depth indentation and, after segmentation, we quantified the distance from the tip at which the membrane deformation reached half of the maximum indentation depth (**Fig S5C**). In control conditions, the distance of the half maximum depth was 1.2±0.2μm and this decreased significantly to 0.8±0.4μm when surface tension was decreased with blebbistatin (**Fig S5D**). Therefore, surface tension participates in setting the length scale of cell surface deformation in response to localised indentation.

The fast vertical displacement is followed by a second slow phase of displacement. With our experimental estimates of k, we can estimate the time-scale of relaxation of a vertical displacement arising from a local deformation of a tensed membrane tethered to a poroelastic cytoplasm. For each position within the region affected by the local deformation either directly or indirectly via surface tension, the time-scale for relaxation is 𝜏_𝑧_∼𝐿^2^/*D_p_*. *D_p_*∼𝑘𝐸 yielding a numerical estimate of *D_p_*∼ 10^-11^ m^2^/s and 𝐿^2^∼𝑙𝛿_𝑧_ with 𝑙∼10 𝜇𝑚 the distance over which vertical displacements are observed and a displacement amplitude 𝛿_𝑧_∼0.5 𝜇𝑚 . With these values, we find 𝜏_𝑧_∼500 ms , qualitatively consistent with the experimentally observed times (**Fig 1D**).

Thus, the instantaneous displacement of the cell surface far from the region of indentation may be due to cellular surface tension (**Fig 1C**) and these vertical displacements may drive intracellular fluid flows throughout the poroelastic cytoplasm.

### Cytoplasmic poroelasticity combined with membrane permeability allows formation of stable intracellular pressure gradients

External stresses applied to the cell surface give rise to transient intracellular pressure gradients that are dissipated by molecular turnover and intracellular flows. However, spatial inhomogeneities in internal stresses generated by the actomyosin cytoskeleton can be maintained for tens of minutes because they arise as a natural consequence of molecular turnover and contractility^34^. For example, during migration, the presence of gradients in myosin motors increasing from front to rear suggests the existence of a high cortical tension at the cell rear that can be sustained without relaxing^10,11^. One implication, given the poroelastic properties of the cytoplasm, is that this surface gradient should result in a sustained pressure gradient continuously driving intracellular fluid flows towards the cell front. Cytoplasmic pressure gradients have been observed in fibroblasts migrating on 2D substrates^14^ and in 3D matrices^9^ ^35^. In addition, intracellular fluid flows have been inferred in cells migrating on 2D substrates^12,14^ and suggested to play a motive role in cells migrating through confined environments^17^.

Because this phenomenon involves a complex interplay between surface tension generated by the cortex, poroelastic behaviour of the cytoplasm, and fluid permeation through the membrane, deriving analytical solutions is challenging and we therefore turned to finite element modelling to gain a qualitative understanding of how long-lasting pressure gradients can be maintained in the cell. In our model, we modelled the cell as a pressurised poroelastic material surrounded by a less permeable surface region 250nm in thickness, representing the membrane and the cortex^29^. For this, we parameterised a poroelastic finite element model (**Fig 4A-B**, **Methods**), adjusting the values of *E* and *D_p_* such that our model could predict the cell’s response to a localised deformation (**Fig 4C**) and the dynamics of displacement of the cell surface in response to microinjection (**Fig 4D-E**).

**Figure 4:**
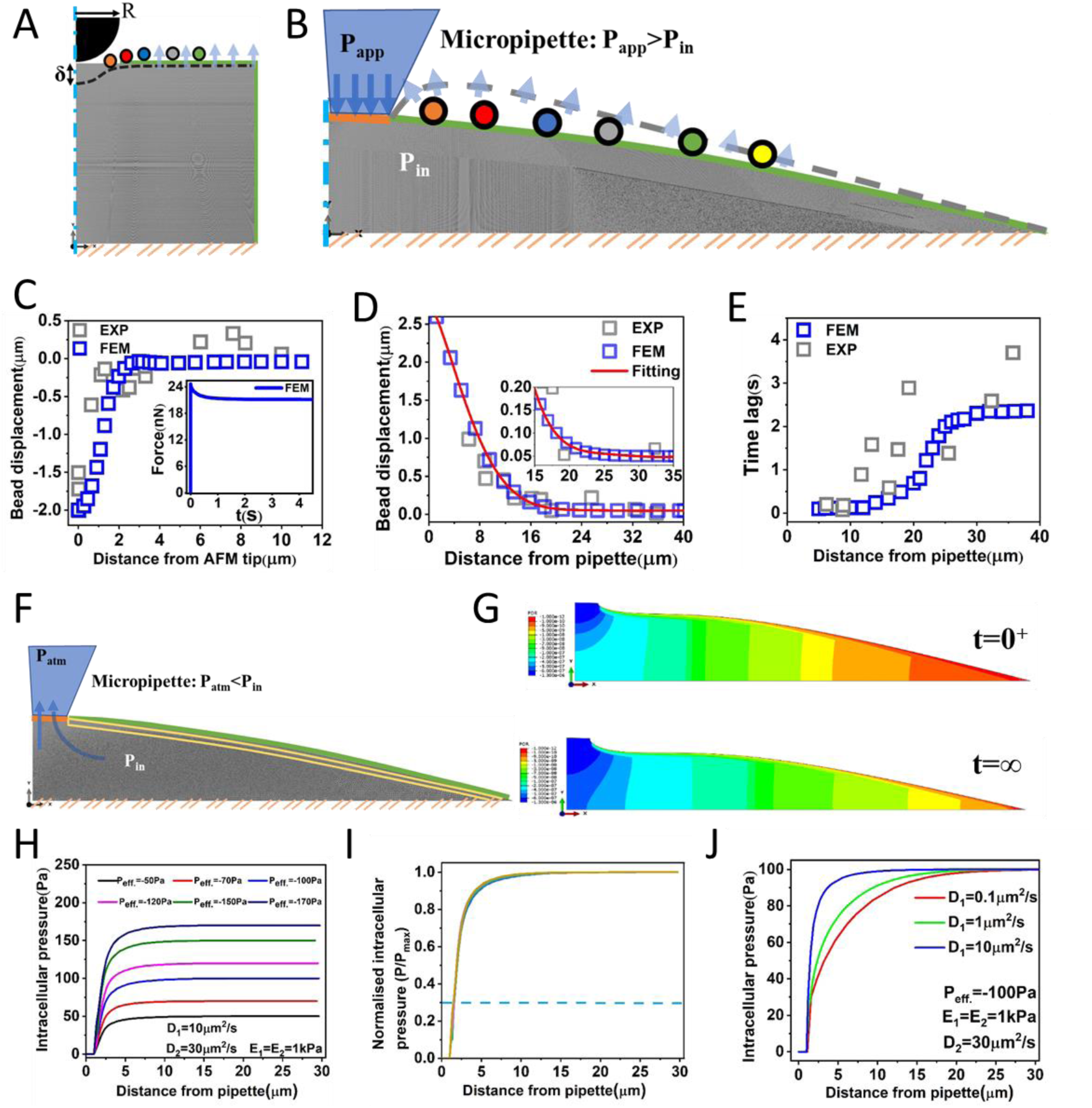
A poroelastic cytoplasm enables the emergence of steady state pressure gradients. **A.** Schematic representation of a quasi two-dimensional slab of poroelastic material with a deformable upper surface. The material is parameterized by its elasticity E, and its poroelastic diffusion constant D. The bottom surface is uniformly tethered to an impermeable infinitely stiff material. The top surface is subjected to an indentation δ applied by a spherical indenter of radius R. The surface profile after deformation is indicated by the dashed black line. **B.** Schematic representation of a two-dimensional slab of poroelastic material (grey). The cytoplasmic material is parameterized by its elasticity E, its poroelastic diffusion constant D, and its pressure P_in_. One part of the surface is permeable and fluid is injected into the cell through this region at a pressure P_app_. The bottom surface is uniformly tethered to an impermeable infinitely stiff material. The surface profile after fluid injection is indicated by the dashed grey line. **C.** Experimental (grey) and predicted steady-state vertical displacement of beads tethered to the cell surface in response to localized indentation. Inset shows the predicted force relaxation as a function of time. **D.** Vertical displacement of beads in response to fluid microinjection after 2s as a function of distance from the micropipette. Grey data points indicate experiments and blue data points the simulation. The red line indicates the trendline. Inset shows a zoom on the further distances. **E.** Time lags of the onset of vertical displacement as a function of distance from the micropipette. Grey data points indicate experiments and blue data points the simulation. **F.** Schematic diagram of depressurization experiment. A two-dimensional slab of poroelastic material (grey) surrounded by a less permeable, thin, outer layer (yellow) representing the cortex. The cytoplasmic material is parameterized by its elasticity E_2_, its poroelastic diffusion constant D_2_, and its pressure P_in._ The outer layer is parameterized by E_1_, D_1_, and P_in_. One part of the surface is permeable and fluid is released from the cell through this region to atmosphere pressure P_atm_. At t=0s, the cell is subjected to a suction Pin through the micropipette. **G.** Pressure distribution immediately after depressurization (top) and at steady state (bottom). **H.** Intracellular pressure as a function of distance from the micropipette for a range of internal pressures P_in_. **I.** Intracellular pressure profiles from H normalised to the pressure at x=30𝜇m from the micropipette. **J.** Intracellular pressure distribution as a function of distance from the micropipette for different membrane-cortex poroelastic diffusion constants D_1_.

After parameterising our model, we then introduced a sink in a region of the cell periphery (2 µm in diameter) to simulate a region of lower pressure (**Fig 4F**). In this region, fluid can rapidly leave the cell, which leads to a reduction in cell volume. In turn, this volume reduction will increase intracellular osmolarity and drive water influx across the membrane in the rest of the cell surface. If the membrane permeability is large enough, efflux through the sink can be compensated by influx through the membrane to maintain a constant cell volume.

When we computationally applied a suction pressure to the sink region, our model predicted a spatial gradient of intracellular pressure with a low pressure close to the sink that increased towards the initial cell pressure far away from this region (**Fig 4G-I**). To gain insights into the importance of the membrane-cortex layer for generation of this pressure gradient, we varied cortical thickness and diffusion constant in our model. While cortex thickness had little influence^3^, diffusion through the cortex strongly affected the length-scale of the gradient (**Fig 4J**), with high diffusion constants leading to gradients that reached the initial cell pressure over short length-scales. This is consistent with previous work that showed that localised exposure of cells to high osmolarity medium leads to localised dehydration with a sharp transition in pore size (and hence *D_p_*) between the exposed and non-exposed region^30^. This work suggested that membrane permeability was on the order of 100 times lower than cytoplasmic permeability, within the lower range of the parameter values tested in our model (red curve, **Fig 4J**). Thus, cytoplasmic poroelasticity combined with passage of water across the membrane can in principle allow maintenance of stable intracellular pressure gradients and pressure compartmentalisation in living cells.

### Cells can accommodate intracellular pressure gradients over minute time-scales

To test our predictions experimentally, we introduced a local depressurisation on the cell surface by establishing a fluidic link using a micropipette and bringing its back-pressure to atmospheric pressure. We verified that, in these conditions, a small suction was generated at the tip of the micropipette (**Fig S6**), signifying that the pressure applied at the pipette tip was lower than the intracellular pressure. Based on the cellular poroelastic properties, the pipette dimensions, and the cell dimensions, steady state is reached for 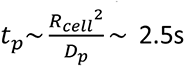 after pressure release, assuming that relaxation is entirely limited by cellular poroelastic properties.

We verified that, following establishment of a fluidic connection between the cell and the micropipette, no occlusion occurred over time and that actomyosin was not perturbed (**Fig S7-8**). To confirm fluid efflux from the cell into the micropipette, we labelled cells with a fluorescent dye that becomes cell-impermeant upon cleavage by cellular proteases. We compared the temporal evolution of fluorescence in pairs of cells, one subjected to depressurisation and a neighbour that was unperturbed (**Fig 5**). Fluorescence intensity remained constant in the control cells but decreased approximately linearly over the course of 10 minutes in the depressurised cells, indicating the presence of a constant pressure gradient and efflux from the cell. Finally, we asked if the cell volume remained constant during depressurisation by examining the change in radius of prometaphase cells subjected to depressurisation. In these cells, cell radius varied by less than 5% over ten minutes of pressure release with no systematic trend to increase or decrease (N=3 cells, **Fig S9**). Overall, these experiments indicate that cells have a sufficiently large membrane permeability to allow rapid exchange of fluid across the membrane to maintain a constant volume and that we can experimentally apply a long-lasting localised depressurisation.

**Figure 5:**
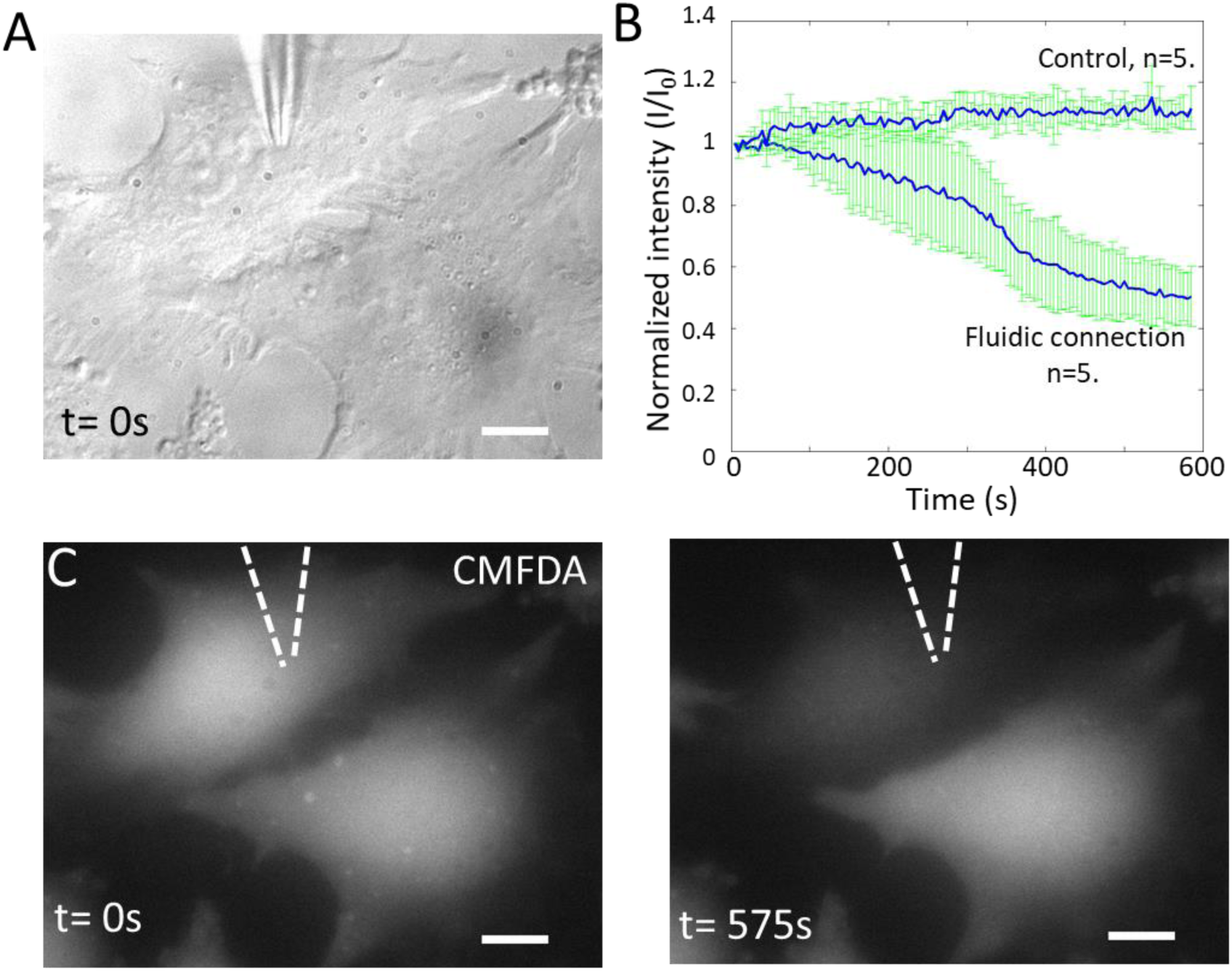
Whole cell patch clamp and pressure release give rise to an outflow of fluid from the cell. **A.** Differential interference contrast image of a typical experiment. The top cell is subjected to pressure release, while the bottom cell is not. Scale bar=5µm. **B.** Temporal evolution of CMFDA cytoplasmic fluorescence intensity in control cells and cells subjected to depressurization. The solid lines indicate the average and the whiskers the standard deviation. N=5 experiments were averaged for each condition. **C.** Fluorescence intensity of CMFDA in the cells in A prior to and after 575s of depressurization. Scale bar=5µm.

Our simulation results indicate that, for values of *D_p_* measured in the cytoplasm, the low pressure region can be confined to a small zone near the pipette if membrane permeability is sufficiently large (**Fig 6C**). To experimentally test this, we used cell blebs as pressure gauges. Blebs are quasi-spherical protrusions of the cell membrane that arise due to pressurisation of the cytoplasm by cortical actomyosin contractility^24,36^. First, we examined naturally occurring blebs in M2 melanoma cells^37^. In control conditions, M2 cells bleb profusely with an intracellular pressure P∼400Pa but, when actomyosin contractility is inhibited with a Rho-kinase inhibitor (Y27632), cells no longer bleb^24^ and intracellular pressure drops to P∼100Pa (**Fig 6A**, see SI methods for pressure measurement). Therefore, if *D_p_* is large and pressure is poorly compartmentalised, local depressurisation should lead to fast global decrease in intracellular pressure and blebbing should cease. In contrast, if *D_p_* is low and pressure is strongly compartmentalised, blebbing should continue unperturbed^38^. In our experiments, following local pressure release (**Fig 6B**), M2 cells blebbed for several minutes, far longer than necessary for intracellular pressure to reach steady state or for significant exchange of fluid to take place (for t>6min, 19/19 cells still blebbed, **Fig 6D**). Furthermore, no spatially localised inhibition of blebbing could be noticed close to the pipette (**Fig S10**). These results suggest that pressure is highly compartmentalised in melanoma blebbing cells and that pressure gradients can be maintained over several minutes. Our pressure measurements indicate that blebs cease to emerge if the intracellular pressure drops below ∼100Pa (**Fig 6A**). Based on this and assuming a diffusion constant of *D_p_* ∼0.01µm^2^/s in the membrane-cortex, our model predicts that pressure-release would only induce a sufficiently large depressurisation to stop blebbing in a region ∼1.5µm away from the pipette tip (**Fig 6C**), consistent with the lack of spatial inhibition of blebbing we observe.

**Figure 6:**
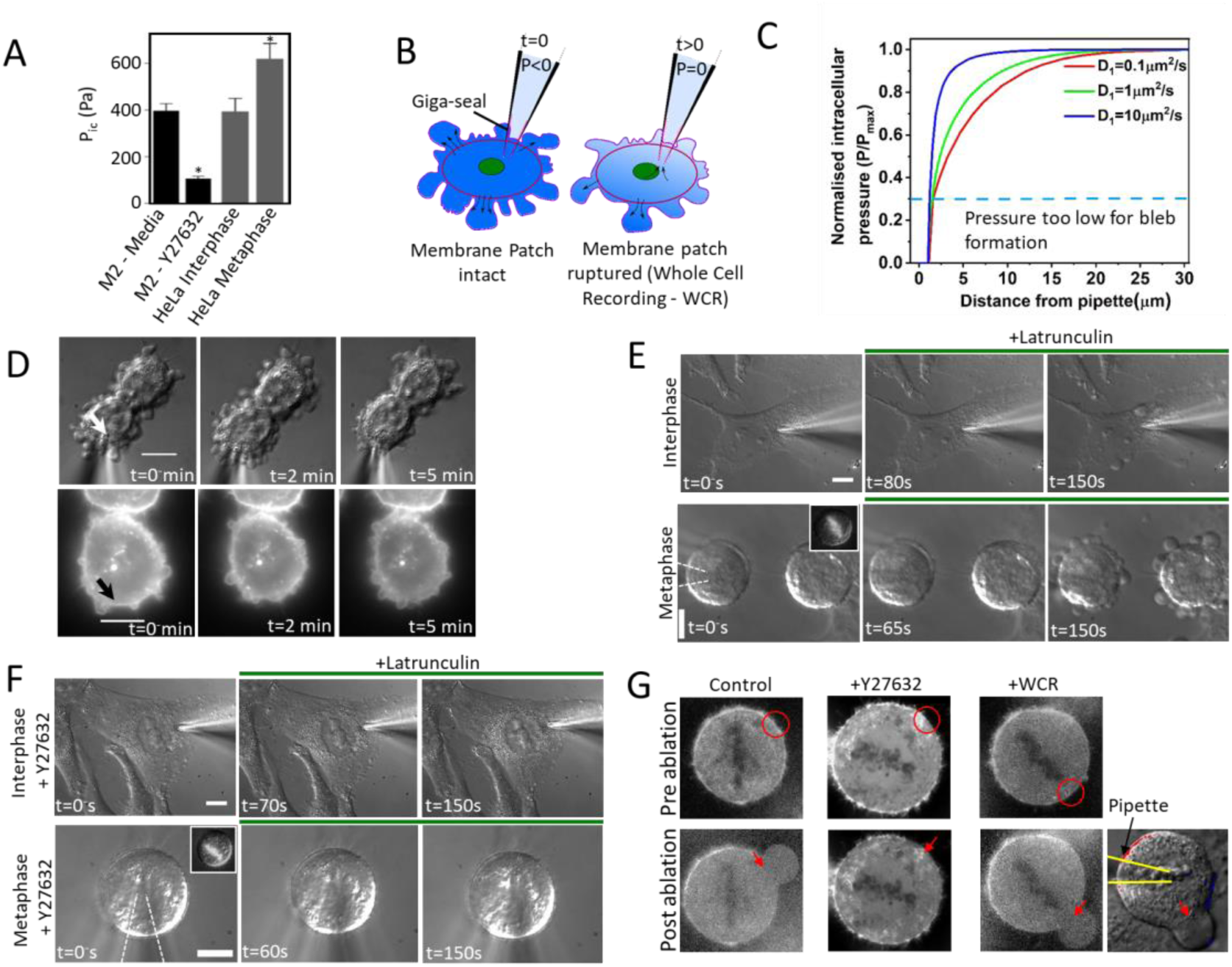
Cells can accommodate sustained intracellular pressure gradients. **A.** Intracellular pressure in Filamin-deficient blebbing M2 melanoma cells and HeLa cells. Cells were treated with Y27632 for 30 minutes prior to measurement. Each condition represents n=34 cells from N=4 experiments. **B.** Schematic diagram of the pressure release experiment. At time t=0s, a fluidic communication is established between a cell and a micropipette, resulting in a small suction pressure at the tip of the pipette. This leads to the establishment of a pressure gradient within the cell. As blebs are pressure driven protrusions, they can be used as pressure gauges to report on the effect of pressure release. **C.** Computational prediction of the intracellular pressure profile in response to depressurization. The dashed line represents the pressure below which blebbing cannot occur based on the pressure measured in control M2 cells and cells treated with the Rho-kinase inhibitor Y27632. **D.** Representative pressure release experiments in M2 cells. Top row: DIC images of a representative experiment. Pressure is released at the pipette tip in a blebbing cell at t=0s and maintained constant thereafter. The location of the pipette tip is indicated by a white arrow. A second cell in the field of view serves as a negative control. Scale bar=10μm. Bottom row: Fluorescence images of the F-actin cytoskeleton in blebbing cells during a pressure release experiment. The location of the micropipette is indicated by the black arrow. Scale bar=10μm. Summary statistics over n=19 cells are presented in Fig S10. **E.** Pressure release experiments in interphase (top) and metaphase (bottom) HeLa cells. Pressure is locally released in the cells at t=0s through a micropipette. The presence of intracellular pressure is detected by the emergence of blebs in response to partial depolymerization of the F-actin cytoskeleton by latrunculin treatment (t=80s and t=150s). Scale bar=10μm. **F.** Pressure release experiments in interphase (top) and metaphase (bottom) HeLa cells treated with the Rho-kinase inhibitor Y27632. (**E-F**) Drugs were added at t=0^+^s. **G.** Laser ablation of the cortex of metaphase HeLa cells expressing GFP-actin. The target region for laser ablation is indicated by the red circle in the before images and by a red arrow in the after images. Control cells, cells treated with the inhibitor of contractility Y27632 for 30 minutes, and cells in which a suction was applied through a pipette are shown. **(E-G)** Summary statistics are presented in **Fig S12B-C**.

We then verified if HeLa cells could also compartmentalise pressure. In these cells, blebs can be induced within ∼2 minutes by partial depolymerisation of the F-actin cytoskeleton induced by treatment with low doses of latrunculin. In these conditions, blebs emerge because HeLa cells have an intracellular pressure ranging from ∼400Pa in interphase to ∼600Pa in metaphase (**Fig 6A**) and because, minutes after latrunculin treatment, cortical actomyosin structures still remain well-defined (**Fig S11**). When contractility is blocked through inhibition of rho-kinase prior to treatment with latrunculin, blebs no longer emerge^39^ (**Fig S11, S12A**). Thus, the growth of latrunculin-induced blebs depends on pressure generated by myosin contractility and they can be used as pressure gauges. When we locally depressurised HeLa cells by establishing a fluidic connection with a micropipette, we found that latrunculin treatment could still induce blebs in both interphase and metaphase cells (**Fig 6E**, **S11B**). This effect was independent of the duration over which depressurisation was maintained. When we established a pressure gradient in cells pretreated with Y27632, we did not observe blebs upon latrunculin treatment (**Fig 6F**, **S11B**).

As latrunculin triggers blebs indirectly and affects the whole of the actomyosin cytoskeleton, we repeated our experiments using blebs triggered by localised laser ablation of the cortex^36,40^. In control conditions, a short pulse of UV laser focused on the cell cortex of a prometaphase cell led to the emergence of a bleb (**Fig 6G**, **S12C**). When myosin contractility was inhibited, ablation did not induce blebs (**Fig 6G**, **S12C**), consistent with ^36^. When a pressure gradient was established, laser ablation could still induce blebs after several minutes (**Fig 6G**, **S12C**), indicating that pressure remained sufficiently large for bleb growth. Collectively, these experiments show that intracellular pressure is strongly compartmentalised in cells and that stable pressure gradients can be sustained over durations of several minutes.

## Discussion

Our experiments revealed that cells could sustain pressure gradients across their cytoplasm over durations of ten minutes, relevant to cell polarisation and migration. In our experiments, we artificially generated an intracellular pressure gradient with a constant high pressure generated by the cell cortex and a low pressure at the tip of a micropipette. Using blebs as reporters of local intracellular pressure, we showed that, despite the presence of an intracellular pressure gradient, pressure over most of the cell periphery remained sufficiently high to continuously generate blebs over several minutes. This result was independent of cell type or cell cycle stage. Computational simulations indicate that the poroelastic properties of the cytoplasm combined with membrane permeability allow the maintenance of stable intracellular pressure gradients.

### Global intracellular flows arise from the combination of cortical tension and a poroelastic cytoplasm

Here we show that local application of stress to the cell surface induces intracellular water flows spatially distributed over tens of microns and lasting seconds. When we locally deformed the cell surface with an AFM cantilever, we observed a global change in cell height with two different temporal regimes: one quasi-instantaneous and another that equilibrated more slowly, consistent with previous work^25^. Previous work has shown that the second phase cannot be explained by a linear viscoelastic behaviour^25^. Therefore, we investigated the role of poroelastic behaviour of the cytoplasm^1^. In line with this idea, when we decreased the

cytoplasmic hydraulic pore size *`ξ`* , the second phase equilibrated slower taking several seconds. Although we cannot exclude a role for complex mechanotransductory processes, our experiments and simulations suggested that the initial fast relaxation is due to simultaneous motion of the solid and fluid phases over a length-scale controlled by surface tension generated by the cortex and the second slower relaxation arise from relative flows between the two phases equilibrating gradients of fluid pressure^26^. However, simple scaling arguments for the poroelastic efflux time based on a hydraulic pore size controlled by homogenous deformation of the solid phase in response to volume change underestimated the magnitude of change in 𝑡_𝑝_, (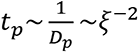 see **Methods**). Indeed, these arguments predicted changes of 1.7-1.9 fold in 𝑡_𝑝_, much smaller than the ∼3.5 fold changes observed in the characteristic equilibration time 𝜏_𝑝_ of the slow phase (**Fig 1D**). This is likely because the hydraulic pore size is governed in a complex manner by interplay between multiple solid structures (cytoskeleton, mitochondria, membrane bounded organelles) and macromolecular crowding.

The existence of spatially distributed intracellular flows induced by local deformation of the cell surface may have important consequences for our understanding of mechanotransduction, which detects mechanical changes and converts them into biochemical signals. Mechanosensory processes act at the molecular scale through opening of ion channels or unfolding of proteins. Thus, the spatial extent of the stress and strain fields will determine what proportion of the cell receives a stimulus sufficiently large and long-lasting to trigger these molecular processes. A challenge for future studies will be to link cellular-scale fluid flows induced by deformation and mechanical stresses to molecular-scale activation of mechanosensory processes to determine the strength of the mechanical stimulus necessary to elicit a whole-cell mechanosensory response. Interestingly, intracellular flows and pressure gradients might participate in triggering signalling through mechanotransductory pathways located away from the cell membrane. For example, Phospholipase A2 activation at the nuclear membrane of epithelial cells is central to the response of tissues to wounding^41^. Thus, the intracellular fluid flows revealed by our study might stimulate intracellular mechanosensory mechanisms, representing an indirect mechanotransductory mechanism.

### Cortical contractility, cytoplasmic poroelastic properties and membrane permeability combine to enable sustained intracellular pressure gradients

Our findings have consequences for our understanding of cell movement in confined environments and low adhesive conditions when cells migrate using blebs^5–7^. At steady state, migrating cells form a stable gradient in myosin that increases from front to rear and that powers migration. At the front, protrusion can either consist of a stable bleb when protrusions grow at a rate similar to actin accumulation^6,7^ or a succession of blebs when actin accumulation rate is faster^5,18^. Disruption of cortical tension at the cell rear by laser ablation leads to cessation of movement as well as localised blebbing in the region of ablation, indicating that myosin accumulation generates a high tension and high pressure at the rear^5^. Our work indicates the cell can sustain long-lasting gradients of pressure and raises the possibility that these may participate in migration. In this picture, the observed gradient in actomyosin distribution might generate an intracellular pressure gradient driving a forward-directed intracellular flow, consistent with some experimental observations^12,13^. In support of this, recent work has shown that new cell protrusions emerge in regions of the leading edge where membrane-cortex attachment is the weakest, suggesting a role for pressure as a driving force^42^. Therefore, pressure gradients may play a direct role in the generation of forward protrusion.

### Conclusions

In summary, our work shows that steady-state gradients in intracellular pressure can arise through the combination of cytoplasmic poroelastic properties and membrane permeability. Thus, intracellular pressure gradients and intracellular fluid flows may play far more important roles than generally appreciated in cell physiology and poroelastic properties must be considered to gain a quantitative understanding of cellular phenomena such as mechanotransduction, cell shape change, and cell migration.

### Author Contributions

MM and GC conceived the experiments. MM performed and analysed all experiments, except the intracellular pressure measurements that were performed by RP and LK. MHE and EM developed the model and performed analysis of the simulations. MM and MHE prepared the figures. GC, MM, MHE, and EM wrote the manuscript.

## Acknowledgements

The authors are grateful to members of the GC lab for feedback and suggestions on the manuscript. The authors wish to acknowledge Prof Guillaume Salbreux and Dr Pragya Srivastava for insightful discussions. MM and GC were supported by grant WT092825 from the Wellcome Trust. GC was supported by a University Research Fellowship from the Royal Society.

## Materials and methods

### Cell Culture

HeLa cells were grown at 37C with 5% CO_2_ in DMEM (Life Technologies, UK) supplemented with 10% FBS (Sigma-Aldrich) and 1% Penicillin/Streptomycin. Melanoma (M2) cells were grown in MEM with Earle’s salts (Life Technologies, UK) supplemented with 10% of an 80:20 mix of newborn calf serum: FBS, and 1% Penicillin/Streptomycin. One day before the experiments cells were trypsinized (Trypsin-EDTA, Life Technologies), transferred from tissue culture flasks into glass bottom tissue culture dishes (Willco Wells, The Netherland). Prior to experiments, the medium was replaced with Leibovitz L-15 (Life Technologies, UK) supplemented with 10% FBS.

To visualise the cytoskeleton, the membrane, and the nucleus, we used previously described stable cell lines: HeLa histone mRFP LifeAct GFP, HeLa LifeAct-Ruby, HeLa MRLC-GFP, HeLa CAAX-GFP and M2 LifeAct-Ruby established using retroviruses and lentiviruses. These were maintained with appropriate selection antibiotics (1mg/mL G418 and/or 250 ng/mL puromycin).

Cells were routinely tested for the presence of mycoplasma using the mycoALERT kit (Lonza). None of the cell lines in this study were found in the database of commonly misidentified cell lines maintained by ICLAC and NCBI Biosample.

### Metaphase arrest

Cells were cultured to reach 70% confluency before being treated with 10 µM s-trityl-L-cysteine (Sigma-Aldrich) overnight to block them in prometaphase. Cells were washed three times using normal medium and then they were released into 20 µM MG132 (Sigma-Aldrich) for 1-2 hours. After this time, many cells were blocked in metaphase. Before the experiments, the medium was replaced with L-15 containing FBS and the same concentration of MG132.

### Microscopy and laser ablation

Differential Interference Contrast and epifluorescence imaging was performed on a Nikon TE2000U (Nikon corp., Japan) inverted microscope. Images were captured on an EMCCD camera (Hamamatsu OrcaER, Hamamatsu, Japan) and transferred to a PC running µmanager (Micromanager, CA). Images were acquired using a 100x oil immersion objective lens (NA=1.3, Nikon) with 2x2 binning. The Ruby fluorophore was imaged using 561 nm excitation and collecting emission at 617 nm. GFP was imaged using 488nm excitation and collecting emission at 515nm.

For some experiments, we used an Olympus IX81 inverted microscope equipped with an Olympus FV-1000 scanning laser confocal head. All images were acquired with a 100x oil immersion objective. Imaging of Ruby and mRFP was performed using a 543 nm laser and imaging of GFP was done using a 488 nm laser.

Laser ablation experiments were performed as described in ^36^ on a scanning confocal microscope (Olympus FV-1000) equipped with two scanning heads. For ablation, the cortex of metaphase HeLa cells was exposed to multiple pulses of a 405nm picosecond pulsed laser (Picoquant). Following induction, blebs grow rapidly before stopping and eventually retracting.

### Chemical treatments

Drug treatments were carried out by adding the appropriate concentration to the medium. Drugs were present at all times during imaging and patch clamping experiments. Latrunculin B (250 nM Sigma-Aldrich) was used to induce blebbing in the cells by inducing partial loss of F-actin. Y27632 (50 µM, Sigma-Aldrich) was used to inhibit actomyosin contractility. Both drugs were dissolved in DMSO. Vehicle controls were carried out by treating the cells with the same amount of DMSO and for the same duration as in the drug treatment cases.

Sucrose (200 mM, Sigma-Aldrich) was used to increase the medium osmolarity, which resulted in shrinkage of the cells and therefore decreasing the mesh size of cytoplasm. EIPA (50 µM, Sigma- Aldrich), an inhibitor of regulatory volume increase (RVI), was used in whole cell patch clamp experiments to prevent volume increase due to transportation of solutes into cell.

### Electrophysiology

The experimental equipment setup consisted of a Digidata 1440A Digitizer and a MultiClamp 700B Amplifier piloted with the pCLAMP 10 Software (all from Molecular Devices, CA). Micropipettes were pulled from thin wall borosilicate capillaires (BF100-78-10, Sutter Instruments, CA) using a Flaming/Brown micropipette puller (Model P-97, Sutter Instruments, CA). Micropipettes had resistance of 6.0 - 6.5 MΩ and a tip diameter of around 2 μm.

For HeLa cells, the pipette was backfilled with a solution was composed of 150 mM K gluconate, 0.005 mM Ca gluconate, 1 mM Mg gluconate, 2 mM K-ATP, 1 mM EGTA, 5mM HEPES, and 5mM Glucose with pH=7.2. For M2 cells, the pipette was backfilled with a solution composed of of 130 mM KCl, 10 mM NaCl, 1 mM MgCl_2_, 5mM Na-ATP, 5 mM EGTA, 10 mM HEPES, and 1 mM CaCl_2_ with pH=7.2. The bath solution was L15 (Gibco Life Technologies, UK) for both cells types and did not contain FBS because this prevents gigaseal formation.

### Data recording and synchronisation

In patch clamp experiments, the pressure transmitter, pinch valves, patch clamp equipment, and the microscope were all connected to the digitizer. This enabled us to control all devices through one platform. The electrophysiology software (pClamp10, Molecular Devices) and the imaging software (Micromanager) were set to communicate to each other. All of the steps to obtain whole cell configuration were performed manually and a macro was created in pClamp 10 to automatically acquire data once whole cell configuration was achieved. At this point by starting the macro in pClamp10, data acquisition, the timing of acquisition of each image, the timing of opening and closing of pinch valves as well as pressure measurement were recorded automatically through one software. This enabled us to synchronize all devices and determine the exact image at which pressure was applied to the micropipette and injection started.

In AFM experiments, for imaging, the camera (Hamamatsu OrcaER, Hamamatsu, Japan) was triggered every 100 ms using one of the output channels of the digitizer. The cantilever displacement and force were also acquired by connecting the corresponding output channels of the AFM to the digitizer using BNC cables. Therefore, similar to patch clamp experiments, we were able to synchronize all equipment by integrating them into one acquisition platform.

### Measurement of intracellular pressure

Direct measurements of intracellular pressure were effected using the 900A micropressure system (World Precision Instruments) according to manufacturer’s instructions and as described in ^9^. Briefly, a 0.5-μm micropipette (World Precision Instruments) was filled with a 1-M KCl solution, placed in a microelectrode holder half-cell (World Precision Instruments), and connected to a pressure source regulated by the 900A system. A calibration chamber (World Precision Instruments) was filled with 0.1 M KCl and connected to the 900A system, and the resistance of each microelectrode was set to zero and then secured in a MPC-325 micromanipulator (Sutter Instrument) within an environmental chamber (37°C and 10% CO 2) on an Axiovert 200M microscope (ZEI SS). To measure intracellular pressure, the microelectrode was driven at a 45° angle into the cytoplasm, maintained in place for ≥5 s before being removed. The pressure measurement was calculated as the mean pressure reading during this interval of time.

### Microinjection and pressure release setup

A pressure sensor (IMPRESS sensors and systems, IMP-LR-C0238-7A4-BAV-00-000) was used to measure the magnitude and temporal evolution of applied pressure. A glass Erlenmeyer was used as a pressure reservoir and was connected to a plastic tube which could be opened by a computer-controlled pinch valve. The tube was then connected to the pressure sensor and the micropipette holder. Two digital manometers were used to monitor the pressure in the reservoir and just before the pipette holder. The length of all tubing was approximately 1 meter and the time for pressure to propagate through the tubes, plus the response time of switches was around 30 ms (data not shown), which is less than our frame interval (66.7 or 100 ms). The pressure transmitter, pinch valves, patch clamp equipment, and the microscope were all connected to the digitizer. Before the start of the experiments, pressure in the reservoir was set. Once the whole-cell patch clamping configuration was achieved, a pulse of pressure was applied by opening the valve, resulting in injection of fluid into the cell. In pressure release experiments, after forming a whole-cell configuration, opening the valve resulted in connecting the pipette to the open atmosphere with pressure of zero (P_atm_ = 0).

### Statistical analysis

All statistical analysis was carried using Microsoft Excel and MATLAB (Mathworks Inc, Cambridge, UK). Custom written codes in MATLAB were used to calculate desired parameters such as: area, speed, time delays, displacements, p value, etc. In all graphs, error bars indicate standard deviation. In box plots, the whiskers represent range of data.

### Image processing

Fiji was used for producing kymographs and preparation of images for the figures.

To extract the profile of the cell surface for **Fig S5**, we cropped the top half of the cell and, for each x position, we determined the z position of the maximum fluorescence intensity closest to the cell interior. Then, to determine the profile of indentation, we subtracted the position during indentation from the one before indentation at each x position.

### Functionalisation of fluorescent beads

Yellow-Green carboxylate-modified fluorescent nanobeads with a diameter of 500nm (FluoSpheres, Molecular Probes, Invitrogen) were coated with collagen-I following the manufacturer’s protocol. To attach nanobeads to the cell membrane prior to experiments, HeLa cells were incubated for ∼30 min with a dilute solution (1:100 dilution) containing the fluorescent collagen coated nanobeads. Unattached beads were then washed out prior to experimentation.

### Defocusing microscopy

Collagen coated fluorescent beads were added to cell culture dishes 30 minutes before the start of the experiments. Some of the beads attached to the cells. The cells were washed in L15 three times to remove unattached beads. Imaging solution was Leibovitz L-15 (Life Technologies, UK) supplemented with 10% FBS. Defocusing microscopy was implemented as described in ^25^. A cell with 2 or 3 attached beads was selected. The motion of integrin-bound fluorescent beads was tracked in three-dimensions using defocusing fluorescence microscopy. By focusing a few microns above the bead plane, each bead appeared as a set of concentric rings. The distance between the beads and image plane is directly related to the radius of the outer ring and is used to determine relative z displacements. Time-lapse images were acquired every 67 or 100 ms and stored as a stack for each bead.

A code was written in MATLAB to automatically track the motion of the beads. Briefly, for a selected bead, a line intensity profile along the diameter of the concentric circles was obtained for each image in the stack. The radius of the outer ring was obtained from the Gaussian fit to the first and last peak in each image. For calibration, changes in radius were then converted into changes Z by moving a bead by a known distance using a piezoelectric stage ^3^ and acquiring images.

### Atomic Force Microscopy and Data Analysis

Indentations of cells by AFM were performed using a JPK NanoWizard-1 AFM (JPK, Berlin, Germany) mounted on an inverted microscope (IX-81, Olympus, Berlin, Germany). The day prior to experimentation, cells were plated onto 35mm glass bottom Petri dishes. Experiments were performed at room temperature and cells were maintained in Leibovitz L15 medium (Life Technologies) supplemented with 10% FBS (Sigma-Aldrich) and MG132 (10µM). Before each experiment, the spring constant of the cantilever was calibrated using the thermal noise method implemented in the AFM software (JPK SPM). The sensitivity of the cantilever was measured from the slope of force-distance curves acquired on glass. For apparent stiffness measurements, we used soft cantilevers with V-shaped tips (BioLever OBL-10, Bruker; nominal spring constant of 0.006 N m^-1^).

For each measurement, the cantilever was first aligned above the cell of interest using the optical microscope. Then, it was lowered towards the cell with an approach speed of 10 µm/s until reaching a force setpoint of 5nN and then kept the cantilever at a constant height.

### Finite element modelling

We conducted finite element simulations to model the mechanical response at the cell scale, specifically focusing on the local deformations of the living cell surface and pressure gradients driving intracellular cytosolic flows. These simulations aimed to capture the deformation behaviour of a poroelastic cell when subjected to changes in effective pressure, mimicking scenarios such as fluid injection or pressure release in one region of the cell surface. Finite element models were developed using ABAQUS (version 2018). We used nonlinear geometry and unstructured mesh in our FE simulations, and also took into consideration a neo-Hookean isotropic porohyperelastic model^3^. This was due to the large mechanical deformation range observed during our AFM, microinjection, and pressure release experiments. The best mesh and domain sizes were determined via mesh convergence studies, and a tolerance for the maximum pore pressure change per increment was calculated for SOILS analysis in our simulations.

#### AFM microindentation simulations-

We ran simulations on two-dimensional rectangular sections to computationally investigate the influence of poroelasticity in the temporal mechanical response of cells to indentation (**Fig 4A**). To minimize edge effects, the cell was represented as a cylindrical disk (20μm radius, 20μm thickness) indented by 2μm with an infinitely rigid indenter representing the AFM cantilever tip. Frictionless and impermeable contact between the indenter and the cell was assumed, and a no-slip condition was imposed on the bottom surface of the cylindre. Pore pressure was set to zero except at the indenter contact surface to simulate fluid drainage. The simulation domain was discretized using quadratic quadrilateral CAX8P elements. Mesh sensitivity checks were performed to ensure independence of results on element size. The simulation consisted of a ramp step followed by a hold step. The ramp step entails indenting the material surface instantaneously (∼0.01s) with specific indentation displacement (∼2um). The phase in which the force reaction is released is known as the hold step. The main parameter of interest was displacement of thr cell surface in response to localized indentation. The force-relaxation curve and final displacement of the cell surface as a function of distance from the AFM tip were analyzed to extract the elastic modulus and poroelastic diffusion coefficient by fitting to the experimental data (For a thorough approach and curve fitting techniques, see ^43^).

#### Fluid injection and pressure release

We ran FE simulations of a poroelastic cell that responds to the application of effective pressure changes at its top boundary where the micropipette contact its surface. The initial cell shape was idealised based on confocal profiles (**Fig 1B**). We modelled the cell as an axisymmetric elliptical cap with a diameter of 40 µm and a thickness of 4.5 µm attached to a substrate. The pipette was in contact with the top surface in the centre of the cell sufficiently long to enable equilibration of effective pressure with the same value as the endogenous cell pressure, *P*_𝑒𝑓𝑓_ = *P*_𝑎𝑝𝑝_*- P*_𝑖𝑛_ = 0. The difference between internal (*P*_𝑖𝑛_) and applied (*P*_𝑎𝑝𝑝_) pressure induces pressure gradients (*P*_𝑒𝑓𝑓_) across the pipette-cell boundary and causes fluid to flow across this boundary. For fluid injection *P*_𝑒𝑓𝑓_ > 0 → *P*_𝑎𝑝𝑝_*> P*_𝑖𝑛_, and for fluid releasing *P*_𝑒𝑓𝑓_ < 0 → *P*_𝑎𝑝𝑝_*< P*_𝑖𝑛_ were considered. **Fig 4B** shows that the pressure increase within the pipette at t=0s led to fluid injection into the cell causing a rise in surface height and pressure changes until it reaches steady state, *P_eff_*=*P*_𝑎𝑝𝑝_*- P*_𝑖𝑛_ = 0. Our simulation had a ramp time (t=2s) over which applied pressure increased the internal pressure. The poroelastic material domain was discretised using the quadratic quadrilateral CPE8P element and the sensitivity of the FE simulations to domain size and mesh element numbers were checked. When the injection is applied, the cell surface reacts with a time lag (*δ_t_*) that increases with the the distance from the micropipette. *δ_t_* is defined as the time for which surface movement reaches 10% of the steady state displacement. In the FE simulations, the elastic modulus—which was extracted from AFM microindentation tests—was taken into account to fit the experimental fluid injection curves and derive *P_eff_* and *D*. To show the impact of the membrane-cortex layer on intracellular pressure gradients and pressure compartmentalisation in living cells, we modelled cells as a double layer poroelastic material^30^. The cytoplasmic material is parameterized by its elasticity E_2_, poroelastic diffusion constant D_2_, and pressure P_in_ and it is surrounded by a less permeable thin layer representing the cortex parameterised by E_1_, D_1_, and P_in_. A no-slip condition was imposed between the two layers.

#### Parameterisation of the Porohyperelastic model -

As in previous studies, we used a fixed Poisson ratio of 𝜐 =0.3 ^3^ and extracted elastic modulus and diffusion constant by fitting experimental steady-state vertical displacement of beads tethered to the cell surface in response to localized indentation with FE simulation (**Fig 4C**). Considering the surface displacement curve, we extracted the following elastic modulus and diffusion constant, *E*=1.8kPa and *D*=28 μm^2^s^-1^, consistent with our experimental results and previous work. To determine effective pressure (P_eff_), elastic modulus (E), and poroelastic diffusion constant (D) for the fluid microinjection of the HeLa cells, we used optimisation processes to fit the vertical displacement of nanobeads positioned on the cell surface after 2s microinjection with FE simulations (**Fig 4D**). The first step in the optimisation process was to conduct the simulations to match the experimental curves and get a first approximations of D and E taking into account P_eff_ =500Pa based on experimental measurements. Next, we adjusted E and D to minimise error compared to the experimental curves and this yielded D= 13 µm^2^.s^-1^ and E=1.2 kPa.

#### Scaling of the poroelastic relaxation time with pore size -

To predict changes in poroelastic relaxation time with cell volume, we tried to gain insight using simple scaling arguments. The poroelastic diffusion constant scales as *D_p_*∼𝜉^2^ , with 𝜉 the hydraulic pore size, and the poroelastic fluid efflux time 𝑡_𝑝_ scales as 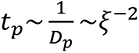. Previous work showed that HeLa cell volume decreases by ∼40% in response to hyperosmotic shock ^30^. The fluid volume fraction 𝑉_𝑓_ in cells is ∼65-75%. If we assume that intracellular water is contained in N pores of volume 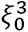, we can express the cell volume as 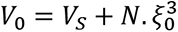 with 𝑉_𝑠_ the volume of the solid fraction. We can rewrite 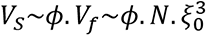 with *ϕ* = [0.42,0.6]. As 𝑉_𝑠_ does not change in response to osmotic shock, we can rewrite the volume change 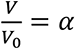 to obtain the change in pore size 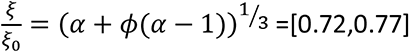 for 𝛼 = 0.4 . This leads to an estimated change in 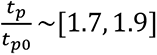

---

**Figure S1:**
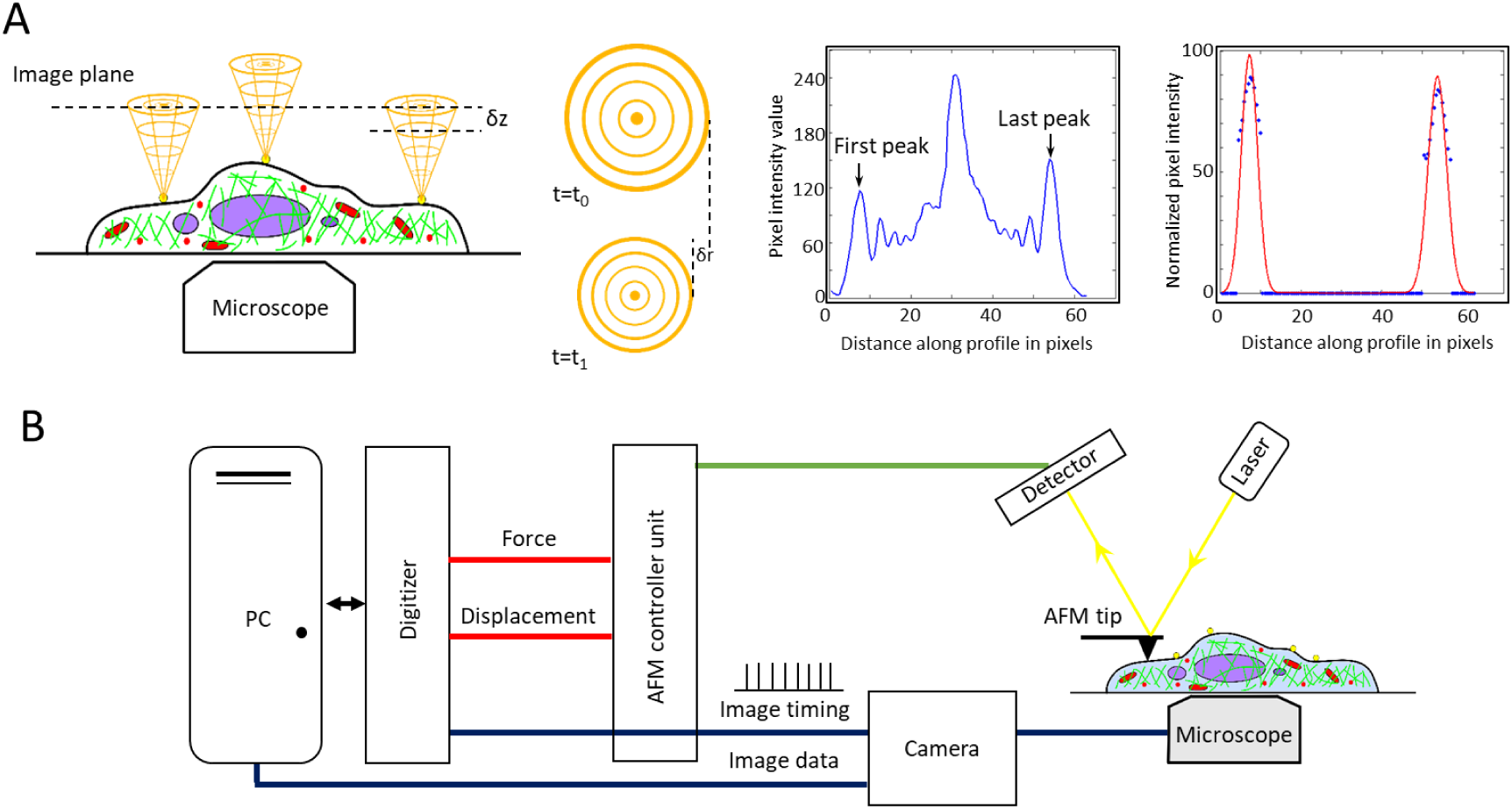
Experimental setup. **A.** Principle of defocusing microscopy. Left: Collagen coated fluorescent beads are bound to the cell surface and are imaged by optical microscopy using a high magnification objective. The plane of focus is purposely set above the cell such that beads appear to have multiple concentric rings around them. The diameter of the outer ring reports on the distance between the bead and the plane of focus. The diameter of the outer ring changes linearly with changes in bead height. The top halo illustrates the rings visualised at t=t_0_ and the bottom halo illustrates them at t=t_1_ once a change in surface height has taken place. Calibration of the outer ring diameter as a function of bead distance from the plane of focus allows the detection of small movements in z^3^. Middle: representative intensity profile of an out-of-focus bead. The central peak denotes the position of the bead centre. Right: detection of outer peak positions. The outer peaks are detected by fitting Gaussian functions (red) to the fluorescence intensity profile of the halo. Blue data points represent the intensity of the outer halo. **B.** Experimental data acquisition setup. An AFM cantilever is positioned above the cell of interest just above the cell membrane. Recording is started and the AFM is brought into contact with the cell to generate an indentation with a depth of 2-4 um. During this time, the position of beads at the surface is monitored using defocusing microscopy. The digitiser simultaneously records the cantilever deflection, the height of the piezoelectric ceramic, and pulses sent by the camera each time an image is acquired. These data allow synchronisation of AFM data and imaging data. Images are recorded separately and are acquired using micromanager.

**Figure S2:**
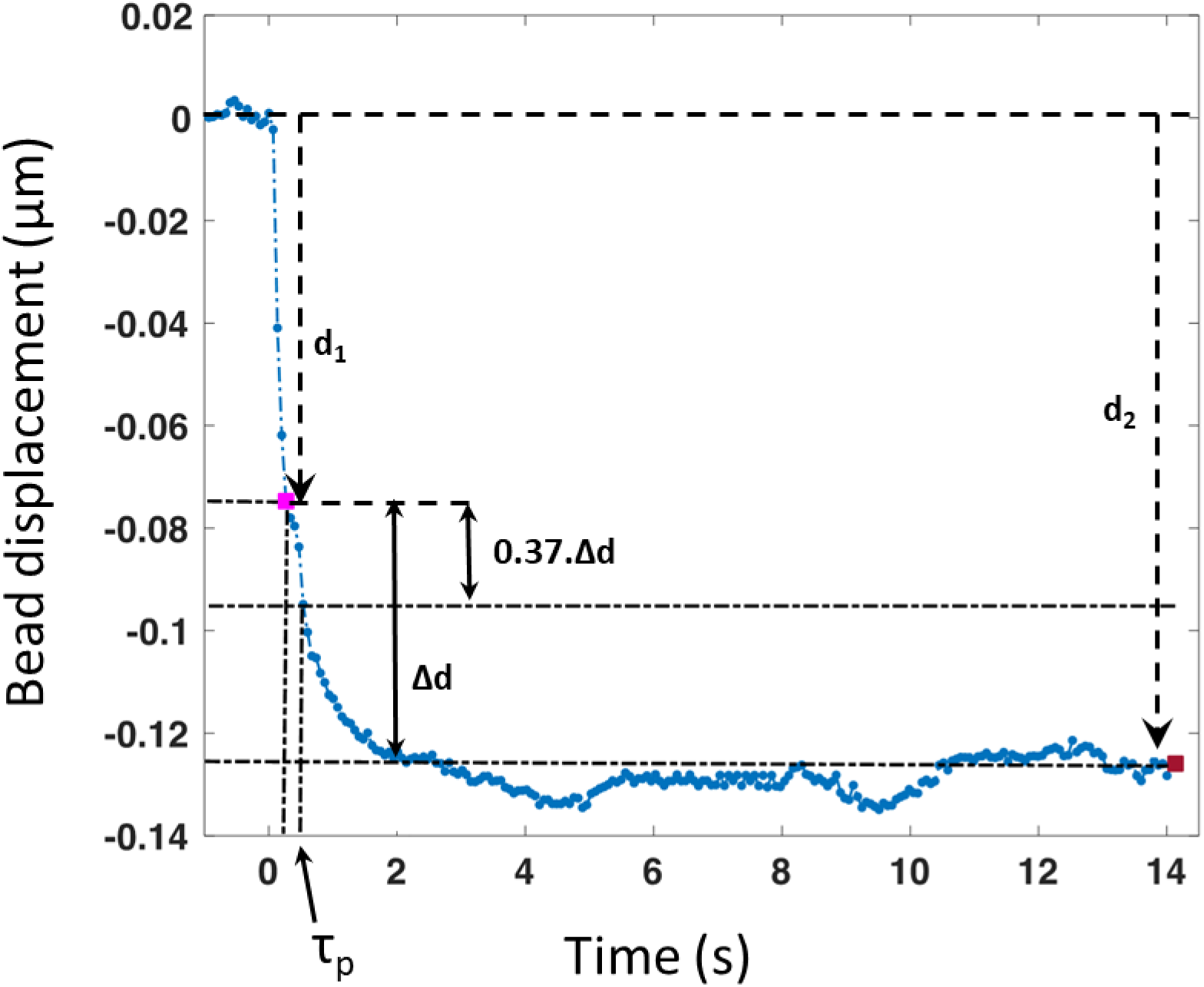
Definition of 𝝉_𝒑_ in AFM experiments. Representative plot of the vertical displacement of a bead tethered to the cell surface as a function of time. The displacement consists of two phases, a first rapid and linear displacement d1 followed by a slower displacement of amplitude Δ𝑑. As previous work has shown that poroelastic relaxation is approximately exponential^1^, we measured the characteristic time 𝜏_𝑝_ . This characteristic time can be defined as the time for which the displacement of the second phase decreases by 37% (1/e) of its initial value: 𝑑(𝑡_1_ + 𝜏_𝑝_) = 𝑑_1_ + 0.37. Δ𝑑.

**Fig S3:**
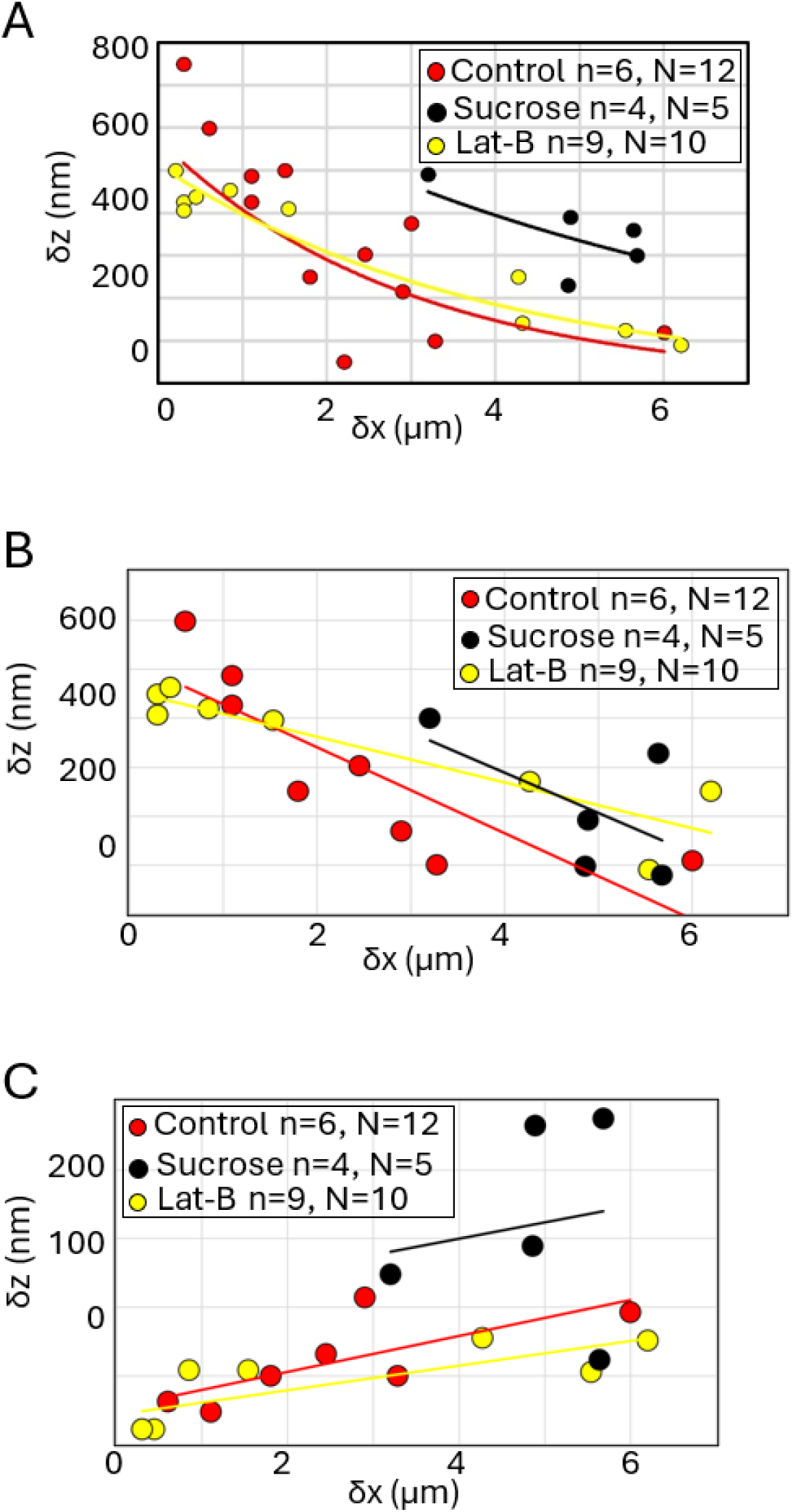
Surface movement in response to application of extrinsic force by AFM indentation. In all panels, experiments in control conditions are shown in red, hyperosmotic conditions in black, and latrunculin treatment in yellow. The number of cells examined (n) and the number of beads examined (N). **A.** Bead displacement δz at steady state as a function of distance to the AFM tip δx. **B.** Bead displacement δz for the first fast phase of displacement. **C.** Bead displacement δz for the second slow phase of displacement.

**Fig S4:**
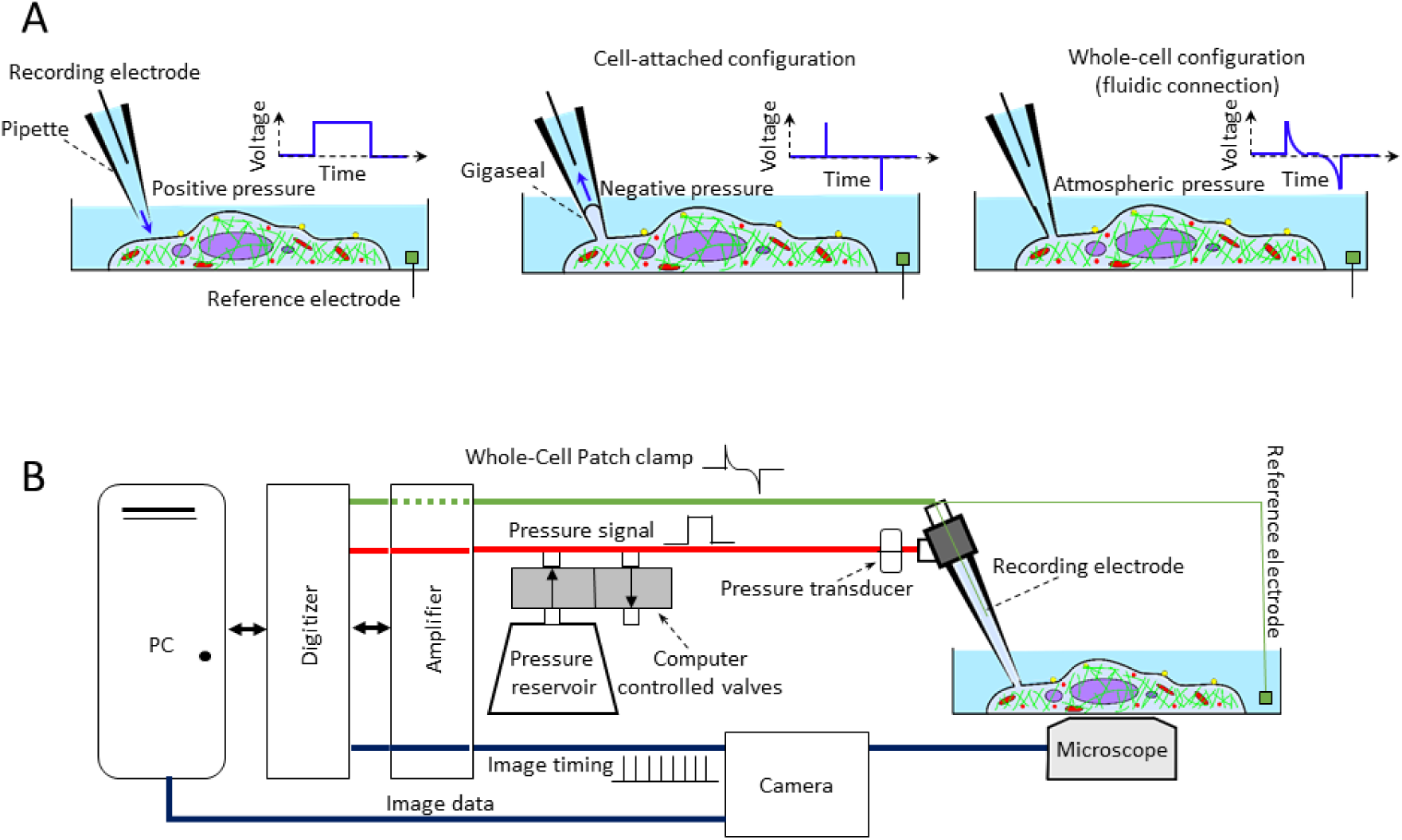
Fluid injection experiments and experimental setup. **A.** Principle of the fluid injection experiments. An electrophysiology setup was used with a whole-cell recording configuration. In patch-clamp electrophysiology, the tip of a micropipette (∼2μm diameter) is first approached towards the cell surface (left) and a high resistance (GΩ) seal is generated by suction of the membrane into the pipette (a configuration known as cell-attached patch clamp, middle). A short pulse of suction can then rupture the membrane patch and create a fluidic and electrical connection between the cell and the micropipette (a configuration known as whole-cell clamp, right). The response to a voltage step can be used to determine the resistance between the two electrodes and reports on the configuration at any given time (indicated as inset in the top right of each panel). **B.** Experimental setup. A cell is brought into fluidic communication with a glass micropipette containing medium with an ionic concentration mimicking intracellular composition. An electrode within the micropipette and within the Petri dish enables measurement of the electrical resistance of the cell-pipette assembly. Pressure can be applied to the fluid within the micropipette using a pressure reservoir and a manifold of computer controlled pinch-valves. Within the micropipette, an electrode is used to monitor the combined resistance of the glass micropipette and cell. Currents recorded by the electrode are amplified before being digitised. The digitiser simultaneously records electrical currents from the electrode in the micropipette, pressure from the pressure transducer, and pulses sent by the camera each time an image is acquired. These data allow synchronisation of electrophysiological data, pressure data, and imaging data. Images are recorded separately and are acquired using micromanager.

**Fig. S5:**
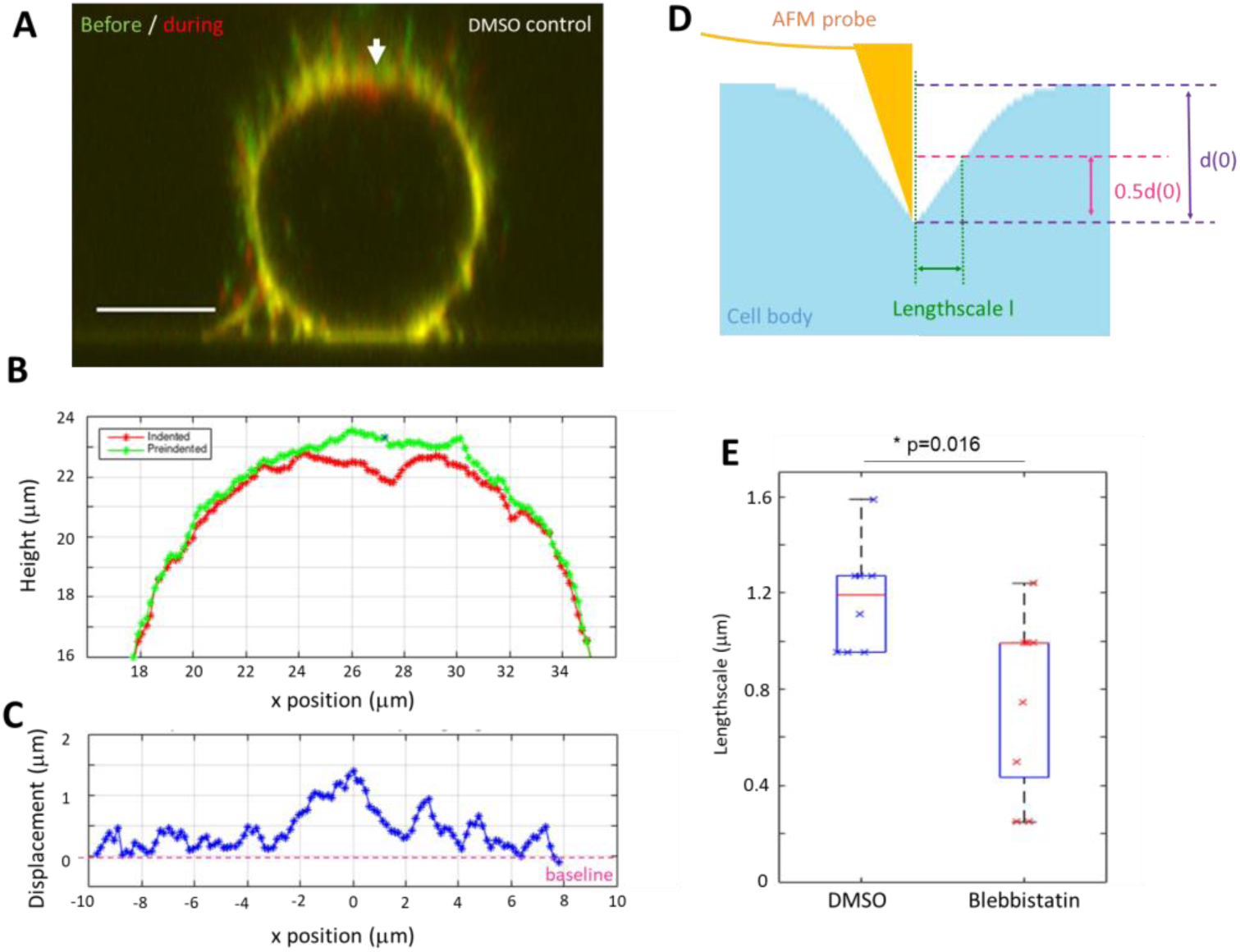
The length-scale of surface deformations is controlled by cell surface tension. **A.** Overlay of the zx profiles of a mitotic cell before (green) and during indentation (red). The cell membrane is labelled with CellMask DeepRed. The arrowhead indicates the position of the AFM tip. Scale bar 10μm. **B.** Segmentation of the membrane along the top half of the cell before (green) and during (red) indentation. Membrane position is derived from segmentation of the data in A. The tip position is marked by an *. **C.** Difference in membrane height between pre-indentation and indentation profiles plotted in B. The tip is located at x=0. **D.** Schematic of the cell surface profile during indentation and the corresponding length-scale of the deformation induced by indentation. **E.** Measured length-scale for an indentation ∼2μm in depth for DMSO control l=1.2±0.2μm (n=8) and for blebbistatin treatment (100μM) l=0.8±0.4μm (n=9) (p= 0.016, student t-test).

**Fig S6:**
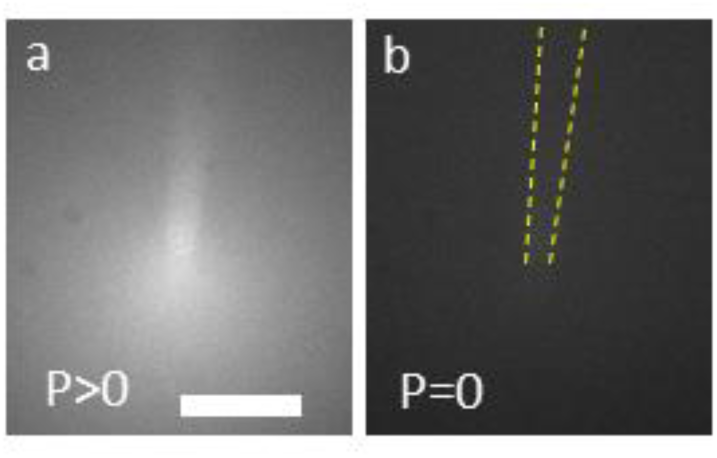
A positive pressure must be applied to the pipette to generate outflow. In all experiments, pipettes were filled with fluid up to mid-height and attached to the micromanipulator at an angle of ∼40°. The pipette medium contained a fluorescent dye and epifluorescence images were acquired by wide-field microscopy. The pipette tip was positioned at a similar height in the Petri dish as during measurements on cells. Scale bar=5µm. **A.** When a sufficiently large positive pressure was applied to the pipette rear, fluorophore leaked out of the pipette. **B.** When no pressure was applied to the rear of the pipette, no fluorophore leaked out of the pipette due to capillary forces resulting from surface tension in the pipette. The position of the pipette is indicated by the yellow dashed lines. The same pipette is shown in A and B. On average, the pressure needed to observe outflow of fluorophore from the micropipette tip was P_c_=0.2kPa (N=5 pipettes).

**Fig S7:**
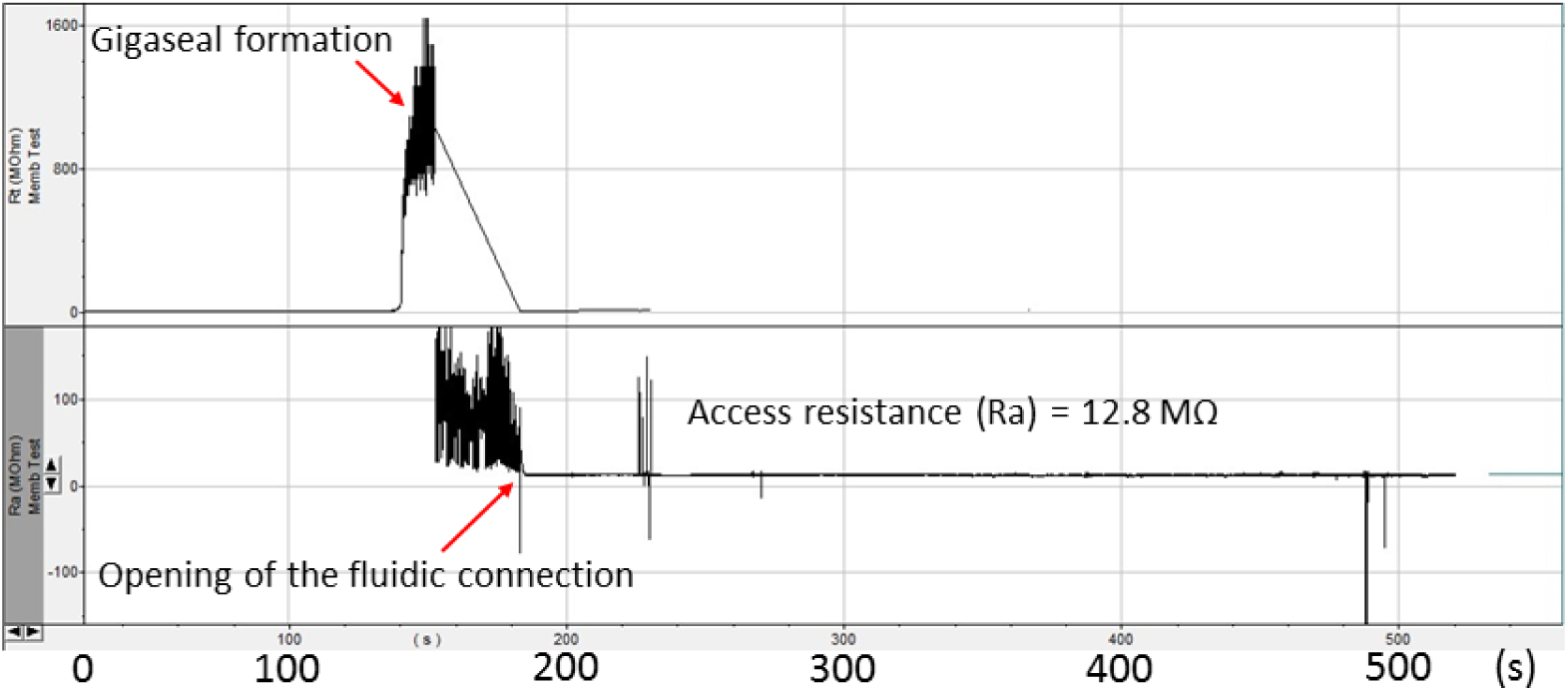
The pipette access resistance stays constant during whole-cell patch clamp configuration. Representative data showing the temporal evolution of the pipette access resistance as a function of time during an experiment in which a suction was applied through the pipette. The vertical axes show the measured access resistance and the horizontal axis shows time in seconds. Two different scales are used in the top and bottom graphs to allow monitoring of gigaseal formation (necessitating GΩ scale, top) and access resistance in whole cell configuration (necessitating tens of MΩ scale, bottom). In the top graph, the micropipette internal fluid is initially just in contact with the cell, a gigaseal is formed by aspiration of a patch of cell membrane into the pipette (see **Fig S4A**), leading to a tight seal between the cell membrane and micropipette interior demonstrated by a large increase of the resistance observed at ∼150s (red arrow, top row). The membrane patch is then ruptured to create a fluidic connection between the cell and the micropipette (while gigaseal is maintained), leading to formation of whole cell configuration with access resistance in the order of Mega Ohms at ∼180s (red arrow, bottom row). In this configuration, a tight seal is maintained between the cell membrane and the micropipette interior. In the bottom graph, the temporal evolution of the access resistance is monitored after formation of the fluidic connection between the cell and the micropipette and during the rest of the experiment. The timing of creation of the fluidic connection is indicated by the red arrow (∼180s). When the fluidic connection is generated, the access resistance (Ra∼13MΩ) is large compared to the pipette resistance (∼6MΩ). This access resistance reports on how easily current can flow between the cell and the micropipette and is extremely sensitive to any clogging or obstacles in the vicinity of the micropipette tip. After generation of the fluidic connection, the access resistance stays constant over the whole duration of the experiment, indicating that a tight seal is maintained between the pipette and the cell and that the fluidic connection does not get progressively clogged by cellular material or active cellular processes. Similar results were obtained for all whole-cell clamp experiments, indicating that no obstruction occurred due to cellular debris.

**Fig S8:**
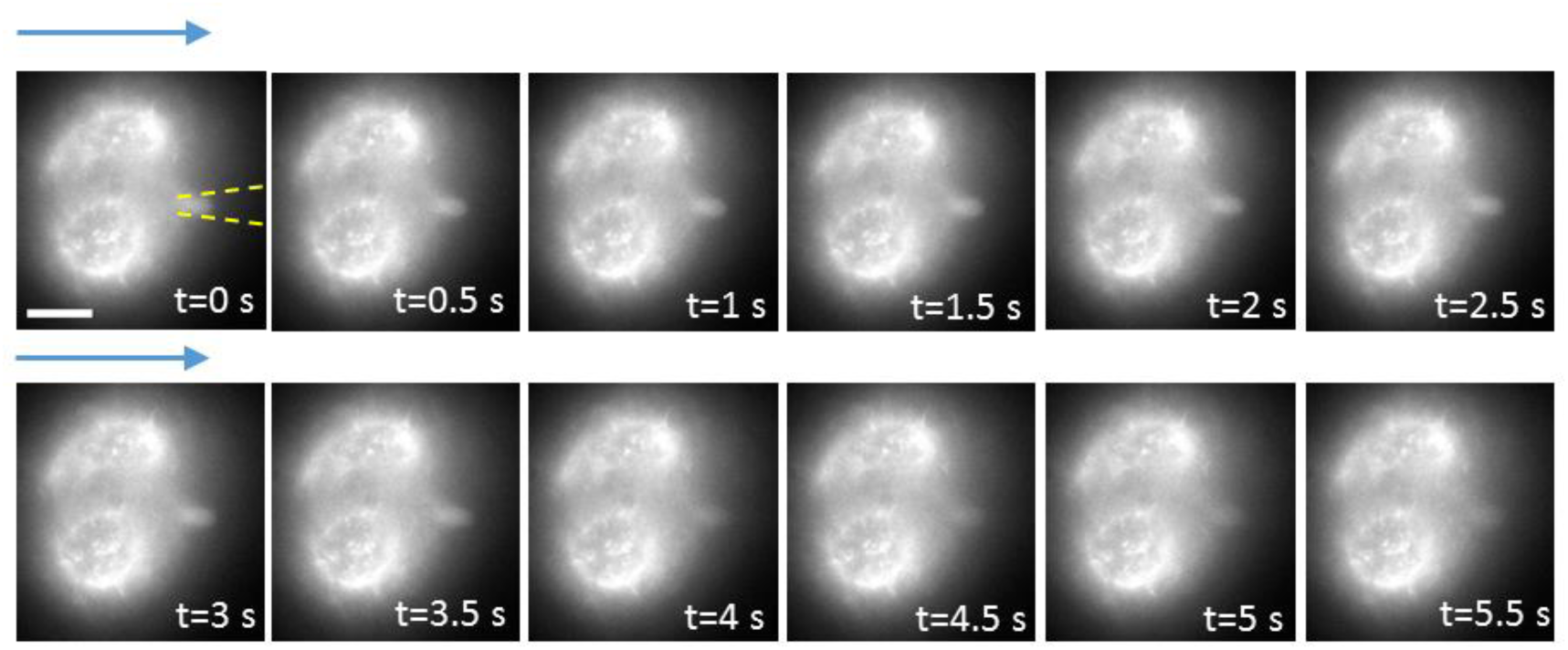
F-actin localisation at the interface between the cell and the micropipette during a pressure release experiment. The cells stably expressed the F-actin reporter Life-Act-Ruby. Pressure release was applied at t=0s through the micropipette. The position of the micropipette is indicated by the yellow dashed lines. All images were acquired by epifluorescence microscopy and show the top of the cell. The distribution of F-actin at the interface between the micropipette and the cell stayed approximately constant over the duration of the pressure release experiment, consistent with the constant access resistance observed (**Fig S7**). Scale bar=10µm.

**Fig S9:**
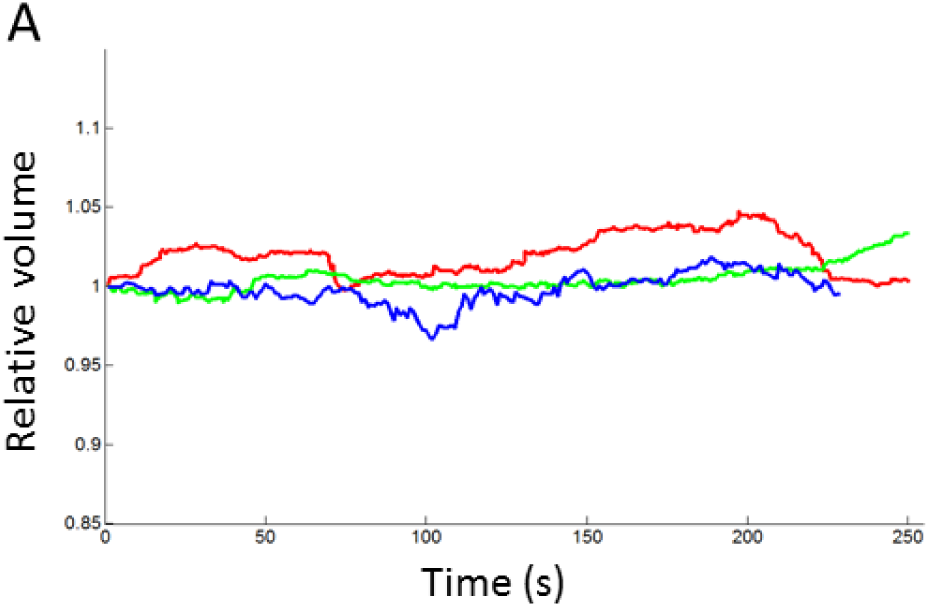
Cell volume remains constant during pressure release experiments. **A.** Temporal evolution of relative volume derived from measurement of the radius R(t)/R(0) for 3 metaphase cells during a pressure release experiment.

**Fig S10:**
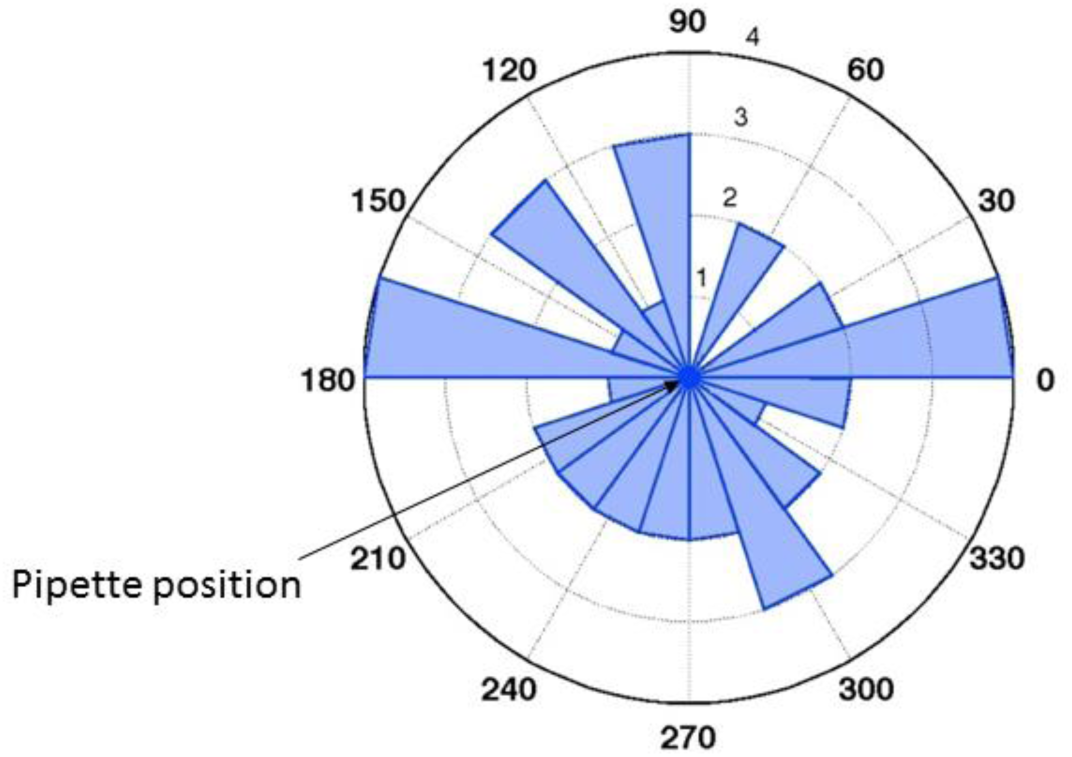
Angular distribution of blebs in a blebbing melanoma cell during a pressure release experiment. Representative data relating to the cell shown in **Fig 6D**, top row. The position of the micropipette is indicated. The concentric circles indicate the number of blebs at each angular position appearing over the duration of the experiments. The experiment lasted a total of 5 minutes. This data is representative of experiments on N=19 cells.

**Fig S11:**
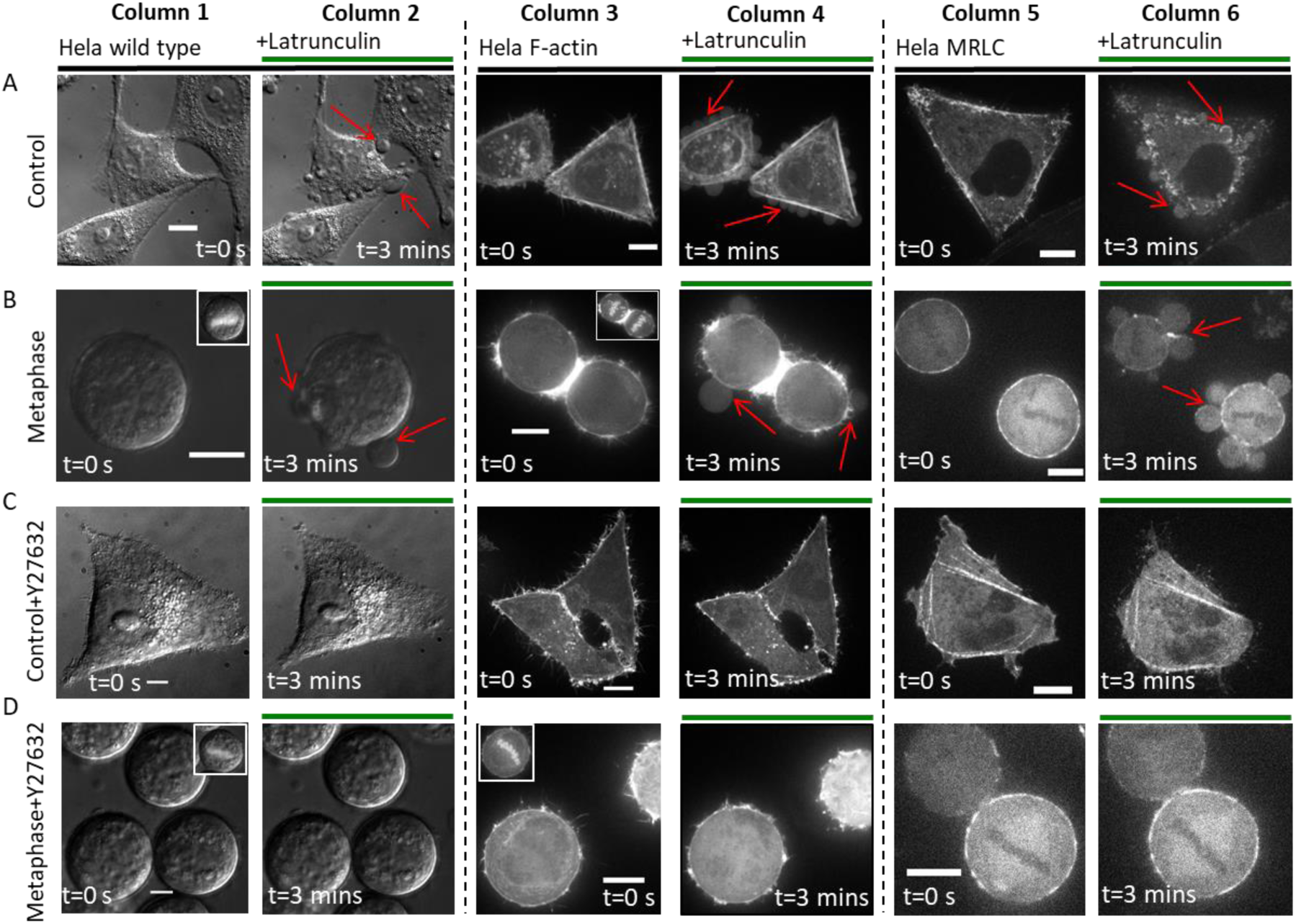
Effect of latrunculin treatment on F-actin and myosin distribution within interphase and metaphase HeLa cells. The first two columns show DIC images of wild-type cells before (column 1) and after Latrunculin treatment (column 2). The 3^rd^ and 4^th^ column show epifluorescence images of cells stably expressing the F-actin reporter Life-act Ruby before (column 3) and after treatment with Latrunculin (column 4). The 5^th^ and 6^th^ column show epifluorescence images of cells stably expressing Myosin Regulatory Light Chain (MRLC) tagged with GFP before (column 5) and after treatment with Latrunculin (column 6). All images were acquired by wide-field microscopy. In all experiments, latrunculin was added at t=0^+^s. In (C-D), cells were pretreated with Y27632 for 30 minutes prior to the start of the experiment. In all panels, blebs are indicated by red arrows. Scale bars= 10um. **A.** Interphase HeLa cells treated with Latrunculin. Treatment with latrunculin gives rise to blebs in all cases but a well defined actomyosin cytoskeleton is still present in the cells after 3 min incubation. **B.** Metaphase HeLa cells treated with Latrunculin. Treatment with latrunculin gives rise to blebs in all cases but a well-defined actomyosin cytoskeleton is still present in the cells after 3 min incubation. **C.** Interphase HeLa cells pretreated with Y27632 to block contractility prior to Latrunculin exposure. No blebs can be observed in response to latrunculin treatment. **D.** Metaphase HeLa cells pretreated with Y27632 to block contractility prior to Latrunculin exposure. No blebs can be observed in response to latrunculin treatment. (**B-D**). Insets show an overlay of the main image with a fluorescent DNA stain.

**Fig S12:**
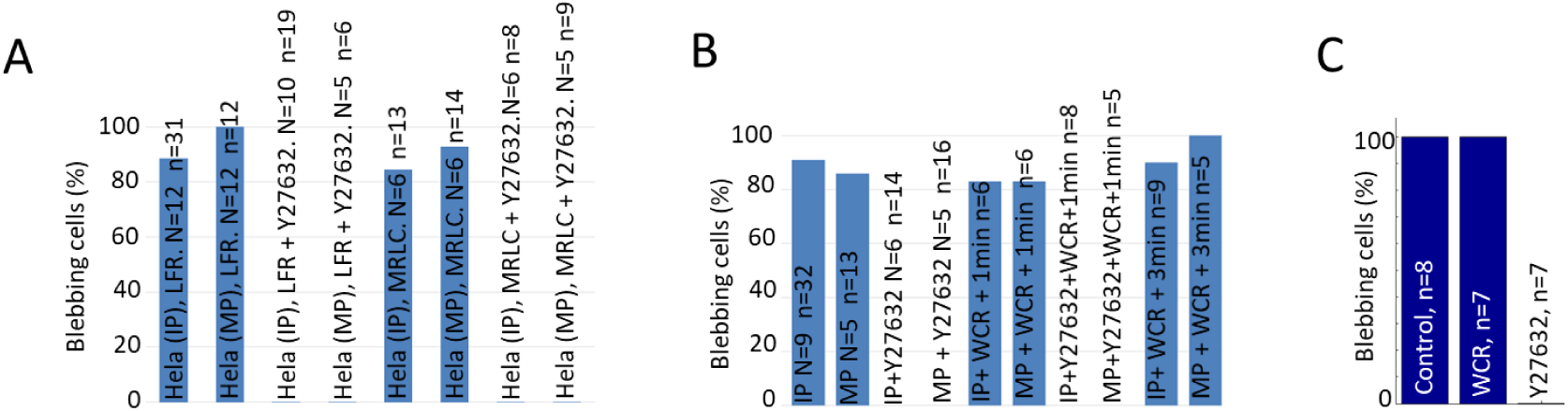
A. Proportion of cells displaying blebs in response to latrunculin treatment with or without pretreatment with Y27632 for 30 minutes. Cells stably expressing Life-Act Ruby (LFR) or Myosin Regulatory Light Chain (MRLC) are used. No differences in response could be observed between LFR cells, MRLC cells, or WT cells (in B). **B.** Percentage of wild-type HeLa cells in interphase (IP) or metaphase (MP) displaying blebs in response to latrunculin treatment for different treatments with (WCR) or without pressure release. Pretreatment with Y27632 was carried out for 30 minutes prior to the start of the experiment. **C.** Proportion of cells displaying a bleb in response to laser ablation for different conditions. WCR indicates pressure release experiments. Pretreatment with Y27632 was carried out for 30 minutes prior to the start of the experiment. On each bar chart, N indicates the number of experimental days and n indicates the number of cells examined.

